# *In silico* generation of synthetic cancer genomes using generative AI

**DOI:** 10.1101/2024.10.17.618896

**Authors:** Ander Díaz-Navarro, Xindi Zhang, Wei Jiao, Bo Wang, Lincoln Stein

**Affiliations:** Ontario Institute for Cancer Research, Toronto, ON, Canada; Department of Molecular Genetics, University of Toronto, Toronto, ON, Canada; Peter Munk Cardiac Centre, University Health Network, Toronto, ON, Canada; Department of Computer Science, University of Toronto, Toronto, ON, Canada; Vector Institute, Toronto, ON, Canada; Department of Medical Biophysics, University of Toronto, Toronto, ON, Canada; Department of Laboratory Medicine and Pathobiology, University of Toronto, Toronto, ON, Canada

## Abstract

Cancer originates from alterations in the genome, and understanding how these changes lead to disease is crucial for achieving the goals of precision oncology. Connecting genomic alterations to health outcomes requires extensive computational analysis using accurate algorithms. Over the years, these algorithms have become increasingly sophisticated, but a severe shortage of open access gold-standard datasets presents a fundamental challenge. Since genomic data is considered personal health information, only an extremely limited number of deeply sequenced legacy cancer genomes can be shared and redistributed. As a result, tool benchmarking is often conducted on the same small set of genomes sequenced with older technologies and uncertain ground truths. This is a major obstacle to the development of improved analytic tools.

To address this issue, we have developed OncoGAN, a novel generative AI tool that uses a combination of generative adversarial networks and tabular variational autoencoders to generate realistic but entirely synthetic cancer genomes based on training sets derived from large-scale genomic projects. Our results demonstrate that this approach accurately reproduces the scale, distribution, and characteristics of somatic point mutations, copy number alterations and structural variants across multiple common cancer types, while protecting donors’ privacy information. OncoGAN accurately recapitulates tumor type-specific mutational signatures as well as the positional distribution of somatic mutations. To evaluate the fidelity of the simulations, we tested the synthetic genomes using DeepTumour, a software capable of identifying tumor types based on mutational patterns, and demonstrated a high level of concordance between the synthetic genome tumor type and DeepTumour’s prediction of the type. We also showed that augmenting real donor data with OncoGAN-generated synthetic data could be used to train a more accurate version of DeepTumour.

This tool will allow the generation of an extensive and realistic set of training and testing cancer genomes whose ground truth is known exactly. This advance provides computational biologists with the ability to develop realistic cancer genome benchmarking sets and make them available to the research community for the testing, development and enhancement of cancer genome analysis tools.

## Background

The vast majority of cancers arise from genomic damage that alters the activity of key genes and proteins involved in the pathways that regulate cell proliferation, programmed cell death, and interaction with surrounding tissues^1^. The promise of precision oncology is that by cataloging these genomic alterations, clinicians can apply therapies that target these altered pathways for enhanced patient outcomes and reduced side effects. Cancer genome alterations have been extensively explored through multiple large-scale genomic projects, starting with the International Cancer Genome Consortium^2^ and The Cancer Genome Atlas^3^, and continuing to this day with such large-scale projects as the PanCancer Analysis of Whole Genomes (PCAWG)^4^, the 100,000 Genomes Project^5^ or the Hartwig Medical Foundation^6^. Collectively, these projects have analyzed over 20,000 whole-genomes from patients with different types of tumors, detecting a large number of point mutations, as well as structural variants (SVs) and copy number alterations (CNAs)^5,7–9^. Using these extensive datasets, researchers have utilized the frequency of these genomic alterations alongside clinical characteristics to identify the key drivers of the cancer phenotype.

To analyze this data, the research community has developed a series of complex computational pipelines, employing multiple variant “callers”^10–14^, each optimized to better detect a specific range of mutations^4^. However, a fundamental issue with these tools is the absence of a gold standard set of cancer genomes with known mutations to test the callers against, and without such a standard it is difficult to know the true error profile of a pipeline, or to rank two pipelines against each other. Computational biologists instead benchmark their mutation calling tools against a small set of approximately 50 legacy genomes derived from cancer cell lines^15,16^, datasets compiled by the Genome In A Bottle Consortium with high-confidence variant calls^17^, or alternatively they test their tools on normal genomes that have been spiked with a known series of *in silico*-generated mutations^18,19^. Unfortunately, none of these approaches fully capture the mutational diversity of actual patient-derived cancer genomes, which vary widely according to the tissue of origin, treatment history, and environmental exposures. Compounding this issue are the legal and ethical restrictions on sharing human genomic data, which make it logistically difficult to assemble and share large numbers of cancer genomes^20–22^. Together these challenges have slowed the development of cancer genome analysis tools.

Recent advances in deep learning techniques such as generative adversarial networks^23^ (GANs) and variational autoencoders^24^ (VAEs), represent an opportunity to create open synthetic datasets that can accelerate the improvement of tumor genome analysis tools. Notably, Yelmen et al. demonstrated the utility of GANs in genomics by constructing a model capable of generating high-quality artificial genomes that simulate haplotype segments across different ethnicities^25^. Briefly, GANs consist of two convolutional neural networks working together. The first one, the generator, creates a result from random data, while the second one, the discriminator or critic, attempts to differentiate real data from synthetic data. By iteratively training both networks, GANs can achieve highly realistic simulations. VAEs, meanwhile, encode input data into a compressed probabilistic latent space before decoding it back. Unlike traditional autoencoders, VAEs introduce randomness by sampling from the latent space, allowing them to generate new, similar data during the decoding process. Although originally developed for image generation, GANs and VAEs have been adapted for tabular data generation through models such as TGAN^26^, CTGAN^27^, CopulaGAN^27^, TableGAN^28^ or CTAB-GAN+^29^ for GAN architectures, as well as TVAE^27^ for variational autoencoder-based approaches.

Here, we present OncoGAN, a pipeline that uses GANs, TVAEs and a random sampling approach trained on large datasets of cancer genome sequencing data to generate unlimited, realistic simulated cancer genomes with known ground truths, including point mutations, CNAs and SVs profiles. The fidelity of these simulated genomes was validated using DeepTumour, a tool that predicts the tumor type of origin based on mutational patterns. Additionally, we demonstrated the utility of these simulations to improve DeepTumour’s accuracy through training on a combined dataset of real and synthetic donors. Importantly, our approach ensures donor privacy, making the simulations fully accessible. Using OncoGAN, we have generated and released 800 simulated genomes across 8 cancer types, which are free of distribution restrictions and ready for use in developing and benchmarking new cancer genome analysis tools.

## Results

### Multi-Model Ensemble Pipeline for Synthetic Tumor Generation

OncoGAN is a comprehensive pipeline designed to generate synthetic tumor samples for eight distinct cancer types (Figure 1). Cancer genomes are characterized by the accumulation of genetic alterations, which can be broadly categorized into ‘driver’ and ‘passenger’ events. Driver mutations directly contribute to cancer development by conferring growth advantages to cells, while passenger mutations are background mutations that are not directly selected. Any individual cancer will have a small number of drivers against a large background of passengers, and a key aspect of cancer genome analysis is to distinguish among these two classes. These mutations can take various forms, including single nucleotide changes, insertions, deletions, and structural rearrangements, all of which contribute to the complexity of cancer genomes. To accurately simulate the complex mutational landscape of point mutations, we trained five different models using the PCAWG^4^, a set of 2,658 whole cancer genomes that have been sequenced and analyzed according to a uniform standard data set (*see Methods*). The models were trained on cancer genome features organized into two primary categories: donor and mutation characteristics. Donor characteristics include: i) the number of different mutation types (single/di/tri nucleotide polymorphisms, SNP/DNP/TNP; small insertions, INS; small deletions, DEL), and ii) the number and type of driver mutations, also referred to here as drivers’ intercorrelation (i.e., driver co-occurrence and mutual exclusion), present in each donor. By training these models independently, we are able to replicate the tumor heterogeneity observed in real data (*see the next section*). Mutation characteristics cover iii) mutational signature contexts, iv) genomic positions, and v) variant allele frequencies (VAF). Since these features are independent and present unique challenges during training, we developed a separate model for each, which were then integrated together.

**Figure 1.**
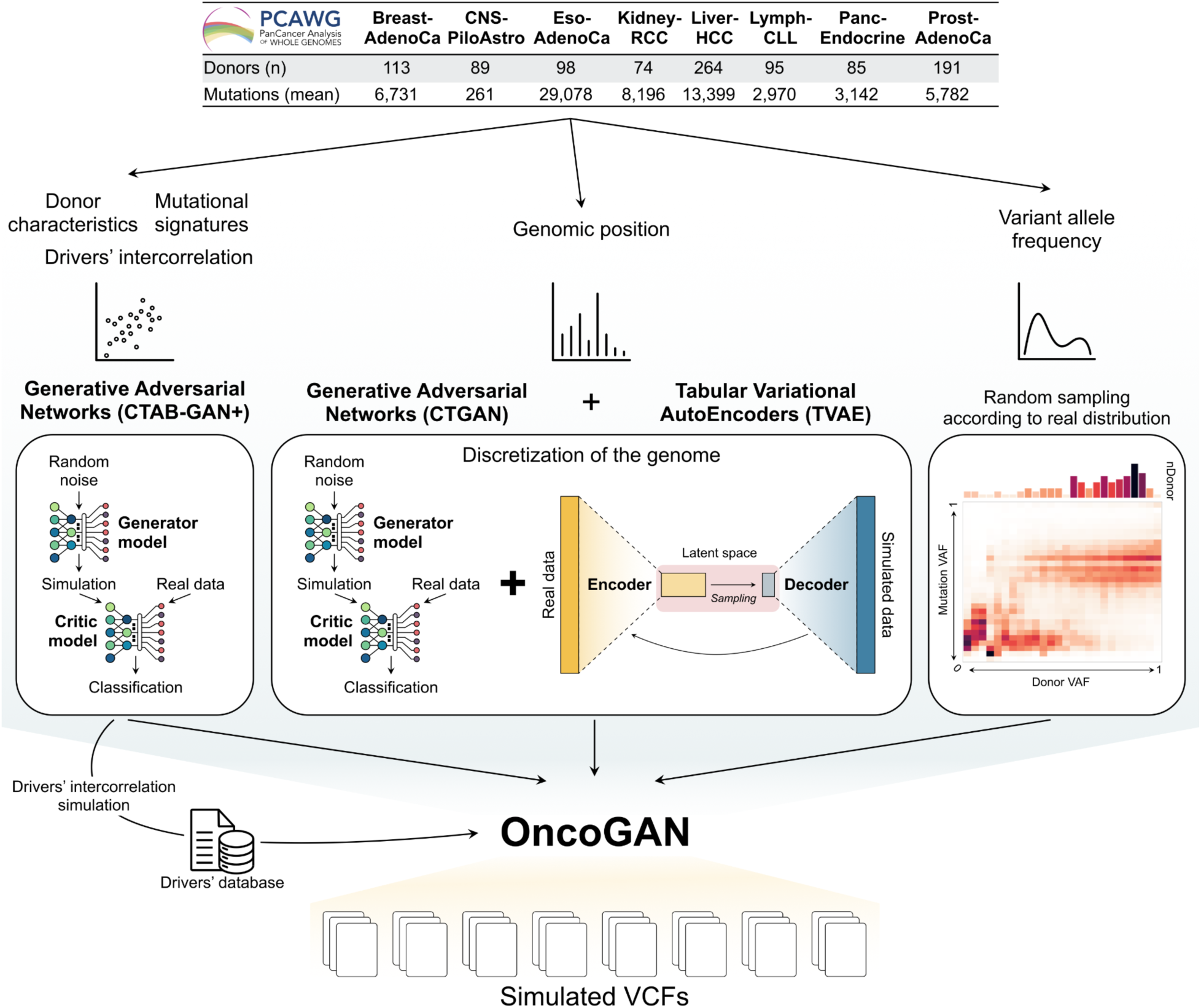
Overview of the OncoGAN ensemble pipeline for the simulation of synthetic VCFs. OncoGAN integrates five distinct models, each trained on data from eight different tumor types sourced from the PCAWG dataset. The architecture of each model, which includes generative adversarial networks (GANs), tabular variational autoencoders (TVAE) and random sampling, is specifically selected based on the input features to ensure accurate simulation. This multi-model ensemble pipeline allows us to generate realistic synthetic donors for the trained tumor types.

We utilized three distinct strategies for training the models: generative adversarial networks (GANs), tabular variational autoencoders (TVAE), and random sampling. CTAB-GAN+ was employed for three of the five models –specifically, the number of each type of mutation and drivers for each donor, and mutational signature contexts– due to their high accuracy in capturing and simulating intra-feature relationships. To predict genomic positions, we combined CTGAN and TVAE architectures with a discretization process (*see Methods*), enabling us to accurately capture mutation density along the genome. Allele frequencies were simulated by first randomly selecting the mean variant allele frequency per donor, based on distributions observed in real patients, and then randomly choosing VAFs according to the expected distribution for that particular mean frequency.

These models were then integrated into an automated pipeline, OncoGAN, which combines their results to generate realistic tumor genomes. The pipeline begins by generating the donor characteristics for the desired cancer type, including determining the number of each mutation type within each sample, the mean variant allele frequency of the donor, and the donor’s sex based on the incidence sex ratio for the chosen cancer type. Subsequently, the number of mutations for each driver gene is simulated, maintaining drivers’ intercorrelation. Since driver mutations cannot be randomly generated, they are selected from a tumor-type-specific list derived from real patient data published previously by the PCAWG^4,30,31^. Following this, the list of passenger mutations, their type, and genomic positions are generated. Given that the models lack reference genome information (i.e., the ability to map coordinates to their DNA sequences), they cannot directly match a generated mutation’s trinucleotide reference context to its position. Consequently, after all mutations are simulated, the trinucleotide contexts and mutation positions are aligned by searching for the reference context within the sequence extracted from the coordinates. A window size of 100 nucleotides around the predicted position is used to search for the expected trinucleotide context, and if no match is found, a new position is simulated to ensure minimal deviation from the initial prediction. This strategy protects donor privacy and ensures that any simulated passenger mutations appearing in the training set do so purely by chance. On average, only 0.021% of simulated mutations exactly match those in the training set, a lower rate than the 0.28% duplication of mutations among donors within each tumor type (Supplementary Table S1).

Finally, a variant allele frequency is assigned to each mutation, resulting in a VCF file for each simulated donor.

Once point mutations are generated, the total number of mutations is used to simulate features of copy number alterations and structural variants. To achieve this, we first trained a CTAB-GAN+ model to simulate i) the number of copy number altered segments and their total length, and ii) the number of inversion and translocation events. Subsequently, three additional models were trained: two CTAB-GAN+ models to independently simulate the desired number of CNA and SV events, and one TVAE model to simulate the positions of SVs, following a similar approach to the one used for point mutations (*see Methods*). The assembly process for CNAs and SVs proceeds as follows. First, aberrant segments are generated based on the number of CNA events generated for a specific donor and tumor type, and their total length is adjusted to match the total simulated length of aberrant copy number alterations. Normal segments are then generated to fill the remaining genome length. Both aberrant and normal segments are shuffled and their sizes adjusted to ensure they remain within chromosomal boundaries. Following the generation of CNAs, deletion and duplication SV breakpoints are automatically assigned based on the copy number profile, ensuring that all segments have their corresponding breakpoints. Finally, other SVs, such as inversions and translocations, are simulated and randomly assigned according to the frequent positions of SVs observed in cancer genomes. SVs that fall within homozygous deletions are excluded.

After generating CNAs and SVs, a plot is produced for each donor, displaying the CN profile alongside the positions of SV events. The CNAs and SVs are saved in the same format used by the PCAWG project.

### OncoGAN Mimics Tumor Heterogeneity and Clonality

To evaluate whether OncoGAN can simulate donors with characteristics that reflect the tumor heterogeneity observed among patients, we generated 100 samples for each of the eight tumor types and compared their characteristics, including the number and types of mutations, their variant allele frequency, and the relationships between driver genes.

#### Mutation density

First, we visually examined the distribution of the number of mutations in both real and simulated donors (Figure 2A and Supplementary Figure S1). This involved comparing the number of each mutation type against the total number of mutations per donor, allowing us to assess not only the range of specific mutations but also their proportion relative to the total mutation count. The resulting scatter and density distributions for the real and simulated data were highly similar, meaning that we are simulating heterogeneous donors. To quantify this similarity, we measured the intersection distance between real and synthetic population histograms for each type of mutation (Figure 2B and Supplementary Figure S2, Supplementary Table S2), where a smaller distance indicates greater similarity. For all simulations, except for certain mutation types in Liver-HCC and Prost-AdenoCa, the results were comparable to those observed in comparisons between subpopulations from the PCAWG dataset, suggesting that the simulated and real populations are indistinguishable. In the case of Liver-HCC, the differences in intersection distances may be attributed to its large sample size, which makes donors with very low or high mutation counts rarer and more challenging to simulate. Nonetheless, the scatter and density plots (Supplementary Figure S1 - Liver-HCC) show that the simulated populations are quite similar. For Prost-AdenoCa, the main difference lies in the simulation of TNPs, likely attributable to the very low frequency of this mutation type, which only represents 0.006% of the total number of mutations, meaning that the differences are minimal. There is also a minor discrepancy in the simulation of deletions. However, the scatter and density plots show no perceptible differences between both populations (Supplementary Figure S1 - Prost-AdenoCa). We also examined the similarity between different tumor types in terms of total mutation counts to determine if OncoGAN simulations exhibit the same patterns as real tumors. To do this, we compared the intersection distances of the different tumors from the PCAWG dataset and our simulations. As shown in Figure 1C, the relationships among tumor types in both cases are nearly identical (Pearson coefficient = 0.966 [0.944, 0.979]).

**Figure 2.**
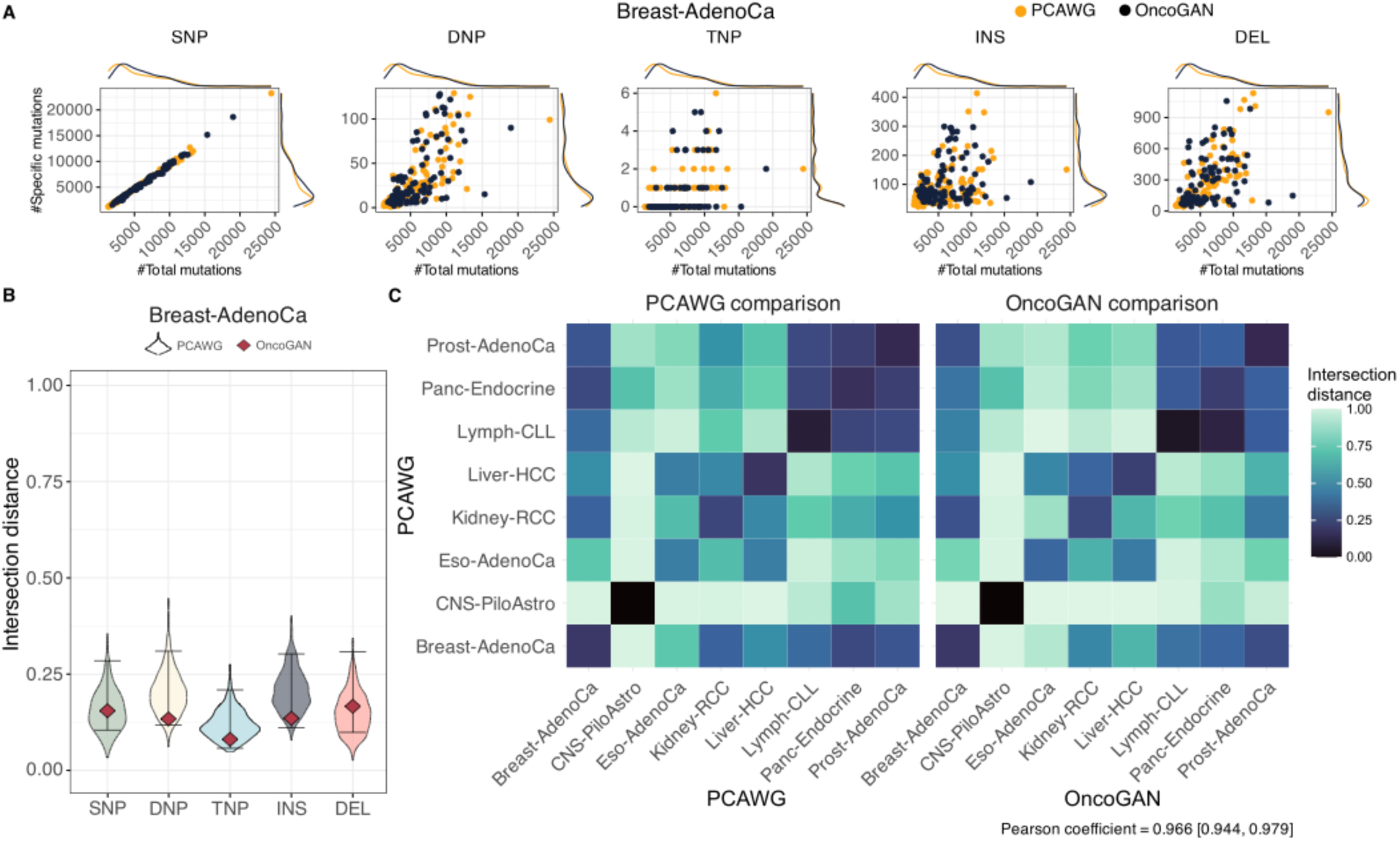
Quality control plots for donor characteristics. A) Scatter and density plots comparing the number of specific mutation types to the total number of mutations for each donor (real in orange, simulated in black) in the Breast-AdenoCa dataset. PCAWG and OncoGAN distributions are highly similar. B) Violin plots for the Breast-AdenoCa tumor type, showing the distribution of intersection distances between two randomly sampled populations from the PCAWG dataset (1000 iterations) and the scores comparing OncoGAN simulations to the actual dataset. The lower the score, the more similar the populations are. For Breast-AdenoCa, all the mutation types have an intersection distance similar to that found in the real data. C) Heatmap of intersection distances comparing different tumors from the PCAWG dataset (left) and OncoGAN simulations (right). A Pearson correlation of 0.966 indicates a high degree of similarity in the relationship between simulated and real donors. A lower intersection distance indicates greater similarity between the two populations. *SNP/DNP/TNP, single/double/triple nucleotide polymorphisms; INS, insertions; DEL, deletions*.

#### Variant allele frequency

Another characteristic simulated by OncoGAN is the donor variant allele frequency, which reflects sample purity and subclonal heterogeneity. This is generated using a simple approach where the donor VAF is sampled from the real distribution, and mutation-specific VAFs are subsequently sampled according to the distribution for that particular donor VAF. As seen in Supplementary Figure S3 and Supplementary Table S3, there are no differences between PCAWG donor VAFs and our simulations, except for CNS-PiloAstro (Wilcoxon test p-value = 0.008). However, the median VAF was 0.222 (q1 = 0.196; q3 = 0.260) and 0.238 (q1 = 0.221; q3 = 0.257) for PCAWG and OncoGAN donors, respectively, with identical variance (F-score = 0.992 and F-test p-value = 0.964 for CNS-PiloAstro), overall indicating that both populations are very similar. When examining VAF for individual mutation types, the distribution is also quite similar between real and simulated data, with two notable differences (Supplementary Figure S4). Tri-nucleotide polymorphisms tend to have a lower VAF in the original dataset, likely due to the small number of TNPs present, while Lymph-CLL indels (insertions and deletions) show a higher overall VAF, possibly resulting from variant callers being less effective at detecting indels at lower frequencies.

#### Driver mutations

The mutations simulated by our models are randomly generated to mimic the background mutational patterns observed in real tumors, and their effects on fitness are not taken into account; hence their frequency distributions reflect those of passenger mutations rather than drivers. To model driver mutations, we sample them from a list specific to each tumor type published by the PCAWG. However, because driver mutations do not appear at the same frequency and some are mutually exclusive, we trained a model to replicate coding and non-coding driver mutation frequencies and their intercorrelation. After this simulation, the correlation between the frequency and inter-relationships of real and simulated driver mutations was very high, with a Pearson coefficient above 0.9 for most of the tumor types (Supplementary Figure S5, Supplementary Table S4). The tumor with the lowest correlation was Eso-AdenoCa (Pearson coefficient = 0.767 [0.759, 0.776]), likely due to the large number of selected drivers for this tumor (10 coding and 60 non-coding), making it more challenging to train an accurate model given the number of available donors.

### Synthetic Tumors Resemble Tissue-Specific Mutational Patterns

#### Genomic distribution

We next sought to confirm that the mutational patterns specific to each type of cancer were accurately generated by our model. While the majority of mutations are passengers, their genomic locations are tumor-type-specific and depend on the cell of origin^7,32,33^. To create high-quality synthetic cancer genomes, it is essential to replicate these specific mutation patterns along the genome. We compared the percentage of mutations found across the genome (in 1Mbp regions) between the PCAWG and OncoGAN datasets, demonstrating that both real and simulated patterns are very similar (Pearson coefficient for Breast-AdenoCa dataset = 0.878 [0.869, 0.886]), with minor differences in some specific regions (Figure 3A, Supplementary Figure S6). A clear example of this accuracy is observed in Lymph-CLL, where we successfully reproduced the mutational peaks associated with somatic hypermutation of the immunoglobulin genes. Moreover, the t-SNE visualization (Figure 3B) shows that PCAWG and OncoGAN donors cluster in a very similar way, easily distinguishing the different tumor types. This suggests that OncoGAN is accurately capturing the genomic positional patterns.

**Figure 3.**
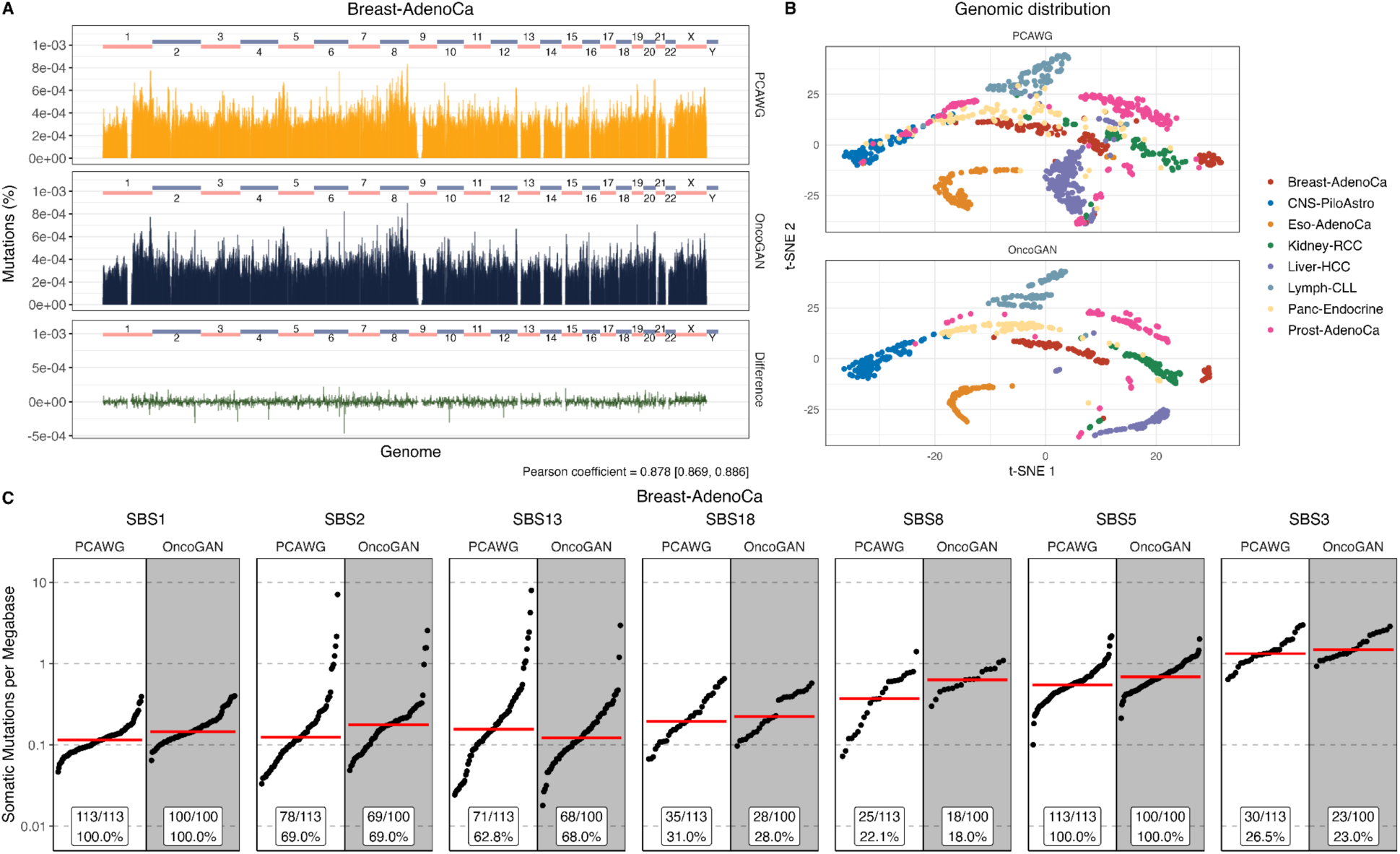
Quality control plots for tissue-specific mutational patterns for Breast-AdenoCa tumors. A) Histogram displaying the total percentage of mutations across the genome in 1Mbp bins. Real and simulated donors are displayed in orange and black, respectively, and the difference is shown in green. A Pearson correlation of 0.878 indicates high similarity between PCAWG and OncoGAN results. B) t-SNE visualization of donors’ genomic distribution (1Mbp windows), colored by tumor type, for both the PCAWG and OncoGAN datasets. C) Distribution of mutation signatures, showing the number of somatic mutations per megabase and the percentage of donors exhibiting each signature. Signatures in the PCAWG dataset are accurately reproduced by OncoGAN, matching both the proportion of affected patients and the mutation density per megabase. Dots represent individual donors; the red line illustrates the mean number of somatic mutations per megabase. *SBS, single base substitutions*.

Similar consistency is also observed when comparing the percentage of mutations detected per region in both datasets (R^2^ for PCAWG = 0.679 [0.638, 0.709]; R^2^ for OncoGAN = 0.772 for Breast-AdenoCa) (Supplementary Figure S7, Supplementary Table S5). The tumor type with the lowest similarity between real and simulated genomic coordinates is CNS-PiloAstro, with a Pearson coefficient of 0.683 and an R² of 0.466. This is comparable to the similarity observed when comparing two samples from the PCAWG dataset, which yields an R² of 0.276 [0.247, 0.301]. These values, which are low in comparison to those of other tumors with a higher mean number of mutations –such as Eso-AdenoCa (R² = 0.977 [0.969, 0.982]), Liver-HCC (R² = 0.983 [0.980, 0.985]), or Kidney-RCC (R² = 0.967 [0.946, 0.982])– can be attributed to the low mutational burden of CNS-PiloAstro (Supplementary Table S5).

#### Mutational signatures

Cancers have distinct mutational signatures related to different biological or external mechanisms, and the mixture of signatures observed in any particular cancer depends on its tumour type, treatment history, and the donor’s environmental exposures. For example, SBS4 is associated with tobacco smoking, SBS7 with ultraviolet light exposure, and SBS9 with the somatic hypermutation activity of the activation-induced cytidine deaminase (AID) in lymphoid cells^7^. To accurately replicate these signatures, we need to simulate the frequency of donors with each signature for a specific tumor type, the density at which each signature is found in each donor, and the relationships among different signatures (e.g., some are mutually exclusive). Thus, we predict the number of mutations for each mutational signature concurrently with the number of specific mutation types.

To test the fidelity of the mutation type distributions produced by OncoGAN, we first compared the distribution of mutation types for each tumour type in synthetic versus real data, demonstrating that our model successfully simulates the inter-tumor heterogeneity among signatures (Supplementary Figure S8). As a further test, we applied SigProfilerExtractor^34^ to reconstruct the mutational signatures in our simulated genomes and compare them to the expected distribution of signatures for each tumour type (Figure 3C, Supplementary Figure S9). We found the SigProfiler mutational distribution profiles derived from the synthetic data to be highly similar to that observed in real donors, with our simulations accurately capturing the variability among donors.

The particular signatures identified by SigProfiler in the OncoGAN-generated genomes matched the signatures identified in real genomes in almost all cases. We were able to identify all expected signatures across six of the eight modeled tumor types. The exceptions were Eso-AdenoCa (SBS10c, SBS8) and Panc-Endocrine (SBS19, SBS30), where two signatures were missing in each type. This limitation is related to the technique we are using. Low-frequency mutational signatures, like SBS10c in Eso-AdenoCa and SBS19 in Panc-Endocrine, are underrepresented in the training dataset (5% of donors) and belong to complex tumor types with eight and ten different mutational signatures, making it more challenging for the model to learn the context distribution among the remaining mutations. In contrast, the SBS30 mutational signature is present in nearly 30% of the original donors. A more in-depth analysis revealed that its primary mutational context (C>T) is being simulated according to its distribution in the training data (Pearson coefficient 0.986[0.96, 0.995]). However, when the training data was compared to the reference profile available in COSMIC and used by SigProfiler, it showed the second-lowest correlation, with a Pearson coefficient of 0.798 [0.515, 0.924]. The lowest correlation was observed for SBS19, which was the other missing signature, with a coefficient of 0.577 [0.038, 0.856]. In contrast, the average correlation for the rest of the detected signatures in Panc-Endocrine was 0.886 (Supplementary Figure S10, Supplementary Table S6).

To further assess the accuracy of the generated signatures, we compared the number of specific mutational signares simulated by OncoGAN with those detected by SigProfiler for each donor. We found that for more than 90% of the donors, these two numbers were highly correlated (Pearson coefficient = from 0.827 (Liver-HCC) to 0.997 (Kidney-RCC)) (Supplementary Figure S11, Supplementary Table S7-S8). OncoGAN simulated a specific signature that was not detected by SigProfiler in only 7.2% of cases. However, in those cases, the simulated signature appeared at a very low frequency. On average, signatures detected by SigProfiler had 2,490 simulated mutations, while those not detected had only 1,342 mutations (Supplementary Table S7). This suggests that these donors may fall below the tool’s detection limit.

#### Indels

Another feature we are simulating, along with the mutational context, is the length of the indels. We compared the indel length distributions in both the PCAWG and OncoGAN datasets and found that they are within the same range, following the same pattern (Supplementary Figure S12). There are some statistical differences for certain indel lengths and tumor types due to sample size, but these do not significantly affect the quality of the simulation.

#### Mutation consequences

Finally, we explored the consequences of the mutations in the genome using the Ensembl Variant Effect Predictor tool^35^, comparing the results between real and simulated donors. As shown in Supplementary Figure S13, for each of the tumor types, the proportion of mutations per donor assigned to the possible consequences are highly similar, with comparable variability across both datasets. Intergenic and intronic variants are the most common categories in all cases, followed by downstream and upstream variants. On average, missense mutations account for 0.64% and 0.66% of the total number of mutations in PCAWG and OncoGAN donors, respectively.

### Performance of Driver Calling Algorithms in Driver Cancer Datasets

As previously explained, the mutations generated randomly by OncoGAN are passenger mutations. However, as we have demonstrated, we introduced driver mutations maintaining the same frequency and inter-relationships observed in real patients. To validate whether the same driver genes can be detected in our synthetic datasets, we applied the widely-used ActiveDriverWGS^36^ algorithm to identify driver genes in both the PCAWG and OncoGAN datasets. The performance of ActiveDriverWGS was highly accurate for both the PCAWG and OncoGAN datasets, detecting 89% and 87%, respectively, of the coding genes for which driver mutations were considered across the eight studied tumor types (Supplementary Table S9). When we examined the driver genes that were not identified by ActiveDriverWGS, we found that these were genes mutated at very low frequencies (<10% donors) in the simulated and real data sets.

### Performance of Tissue of Origin Prediction

Ensuring that simulated genomes not only replicate individual characteristics but also maintain tumor-type specificity is essential for validating their biological relevance. As we have shown previously, the individual characteristics OncoGAN simulates closely resemble the original data. To assess whether the final simulation (i.e., each individual simulated donor) corresponds to the expected tumor type, we utilized DeepTumour, a tool that predicts the tumor tissue of origin based on the list of detected somatic mutations^37^. We ran DeepTumour on the synthetic donors and achieved very high prediction accuracy, nearly 100% for most of the tumor types (Supplementary Table S10). However, when examining the 5-fold cross-validation results of the original model, we observed that some tumor types had lower performance (Figure 4C - Baseline metrics). In particular, we investigated Lymph-MCLL and Eso-AdenoCa datasets in more detail to understand the drop in performance for these tumor types.

**Figure 4.**
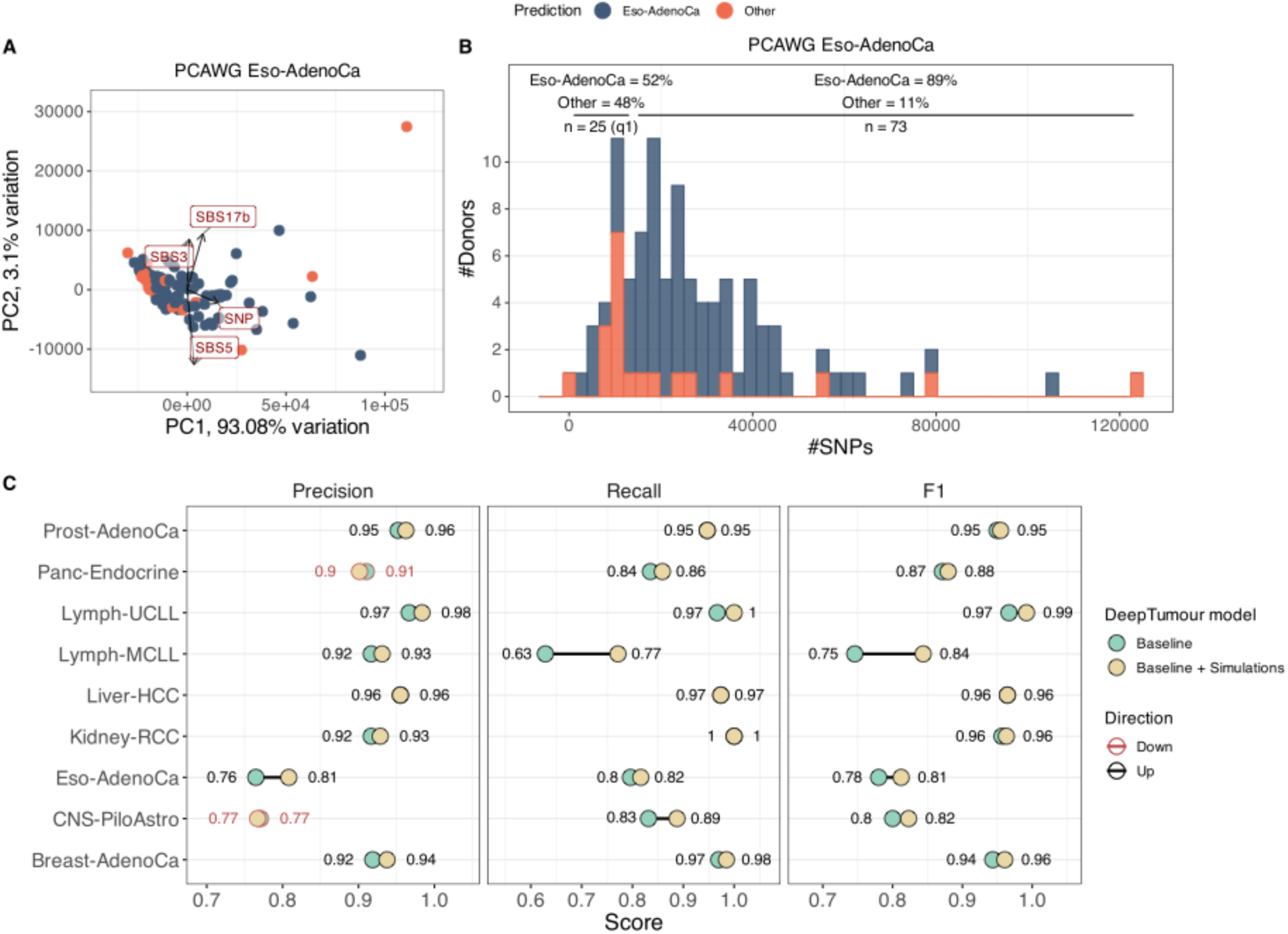
DeepTumour prediction analysis. A) PCA of Eso-AdenoCa real donors using the total number of each mutation type and mutational signatures as features. The four most significant features are highlighted in red boxes. B) Histogram showing the distribution of accurate and misclassified Eso-AdenoCa donors by DeepTumour according to the number of SNPs. Donors with fewer mutations are more frequently misclassified (48% vs. 11%). C) Comparison of precision, recall, and F1 metrics between the baseline DeepTumour model (light green) and a model trained with the original dataset plus 100 simulated donors for each of the eight tumor types (beige). Metrics that decreased in the new model compared to the baseline are marked in red text.

Lymph-CLL is a disease characterized by two distinct subtypes, U-CLL and M-CLL, based on the mutational status of the immunoglobulin heavy-chain variable region (IGHV), with M-CLL presenting three mutation peaks at the IGHV genes (chr2-IGK, chr14-IGH, and chr22-IGL, Supplementary Figure S6) and the mutational signature SBS9, while U-CLL lacks these features37. When analyzing the 5-fold cross-validation results from DeepTumour, we observed that the recall for Lymph-MCLL was the lowest, with only 63% (22/35) of donors correctly classified, and the remaining 37% (13/35) misclassified as B-cell non-Hodgkin lymphoma (Lymph-BNHL) (Supplementary Table S10). This misclassification can be attributed to the nearly identical mutational patterns between these two tumors, with a Pearson coefficient of 0.941 and R^2^ of 0.887 (Supplementary Figure S14). The primary difference occurs in the three regions that correspond with the IGHV genes, which have a lower percentage of mutations, as only 73% of Lymph-BNHL cases present the SBS9 signature, compared to 100% of M-CLL donors. Furthermore, of the 95 Lymph-CLL donors used to train DeepTumour, only 35 (37%) were M-CLL, indicating that a larger sample size may be necessary to improve the prediction accuracy for this tumor type.

On the other hand, Eso-AdenoCa had the lowest precision (76%) and the second lowest recall (80%) among the tumor types. To investigate the Eso-AdenoCa donors, we performed a PCA analysis based on the number of each type of mutation and signature. As shown in Figure 4A, misclassified donors can be mainly distinguished from the rest by their number of SNPs. Specifically, when dividing the donors by their total number of SNPs, we found that those with the fewest mutations (Q1, n=25) were more frequently misclassified (48% vs. 11%) (Figure 4B). This suggests that these samples may represent cases that are underrepresented in the training dataset.

### Improving Algorithms Performance Combining Real and Synthetic Datasets

Given these findings, we decided to train a new DeepTumour model using a mixed dataset of real and synthetic donors to determine whether this could enhance the model’s accuracy. We trained DeepTumour with the same 5-fold cross-validation strategy, but adding 100 OncoGAN-simulated donors to 8 of 29 tumor types present in the training dataset. For this new model, the overall accuracy improved by 0.9% (from 89.26% to 90,16%) (Supplementary Figure S15). For the eight studied tumor types, the 5-fold cross-validation precision, recall, and F1 scores increased by 1%, 1.58% and 1.34%, respectively. Additionally, tumor types with fewer donors in the original DeepTumour training dataset benefited the most from the supplementary training, such as Lymph-MCLL (n=35) and Lymph-UCLL (n=60), with F1 score increases from 75% to 84% and from 97% to 99%, respectively (Figure 4C). On the other hand, two tumor types experienced a drop in one of the metrics: CNS-PiloAstro (−0.4% precision) and Panc-Endocrine (−0.9% precision), likely due to an increase in Thy-AdenoCa donors misclassified as CNS-PiloAstro, and CNS-Oligo donors misclassified as Panc-Endocrine, possibly due to the low number of donors available for those tumor types (n=48 and n=18, respectively) (Supplementary Figure S15).

### Expanding the Simulation to Copy Number Alterations and Structural Variants

To simulate an accurate representation of copy number alterations and structural variants for each donor, we trained a model that uses the total number of point mutations as the basis for generating these alterations (Supplementary Figure S16). For CNAs, we define segments of varying lengths and simulate the minimum and maximum number of alleles for each segment, ensuring these features follow the same distributions observed in real patients. To measure the quality of the simulations we calculated several chromosomal instability scores (CIS), such as the total number of segments and their mean size, the fraction of the genome that is altered (FGA) or the total aberration index (TAI) (Figure 5)^38^. The simulated tumor types show high concordance with the PCAWG dataset, with the exception of Eso-AdenoCa, which differs in the FGA metric due to the simulation of a higher number of altered segments. For SVs, duplication and deletion events were automatically assigned based on copy number status, while the number of inversions and translocations was simulated to match the distributions observed in real donors. As shown in Supplementary Figure S17, the number and lengths of these events between OncoGAN and PCAWG donors for each tumor type are highly similar. In the case of translocations, PCAWG does not report the actual size of each event, since for most translocations only a single breakpoint is detected. Although the comparison between real and simulated donors is statistically significant in many cases (Wilcoxon test p-value < 0.05), the differences between the two datasets for the eight different tumor types are small, as evidenced by the similarity of their means (Supplementary Table S11).

**Figure 5.**
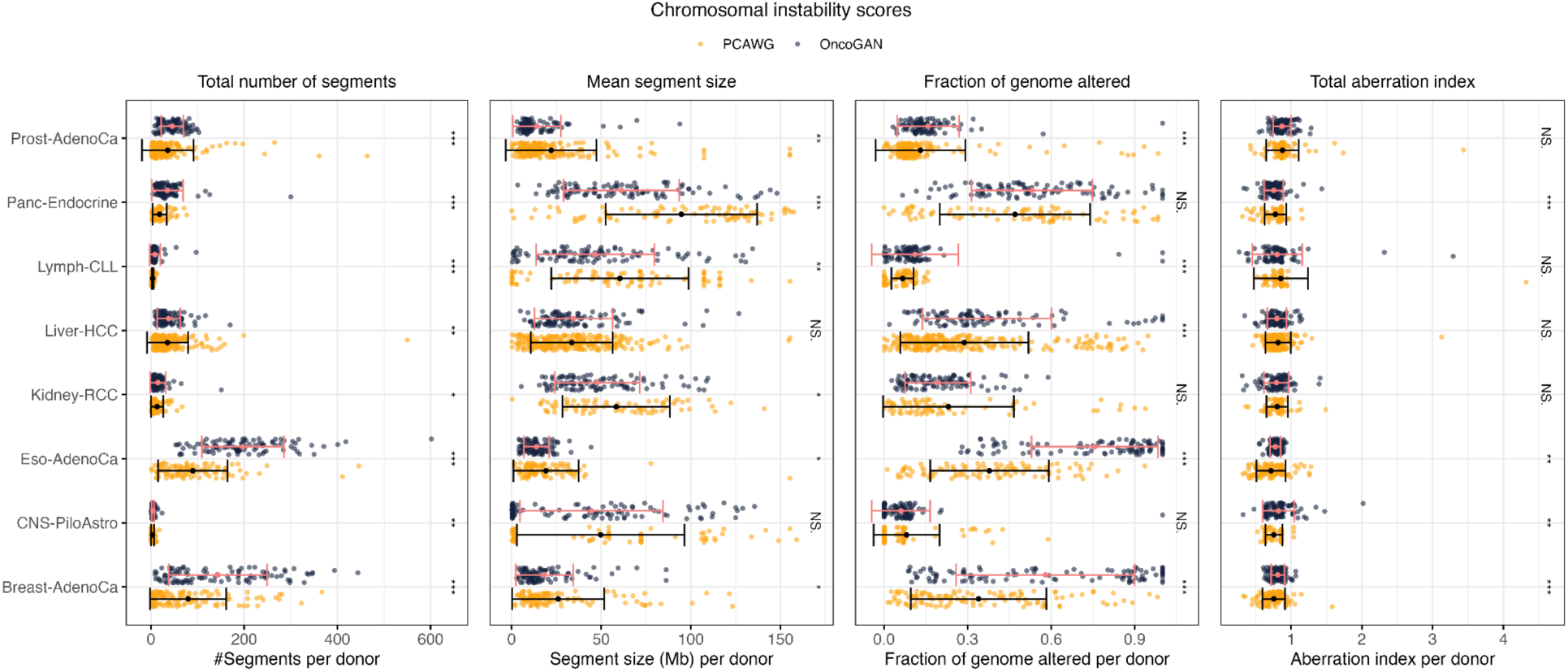
Chromosomal instability scores to measure the similarity in copy number alterations between real (orange) and simulated (black) donors, demonstrating high concordance between the two groups. The total number of segments indicates the number of altered regions per donor (i.e., regions where the major and minor alleles differ from one). The mean segment size refers to the average length of the altered regions. The fraction of the genome altered is calculated by summing the lengths of all altered segments and normalizing by the total genome size. The total aberration index (TAI) measures the average deviation from the normal copy number state. Error bars show the standard deviation of the dataset. The Wilcoxon test was used to compare the groups. *NS.: p-value > 0.05; *: p-value <= 0.05; **: p-value <= 0.01; ***: p-value <= 0.001*.

### Compatibility with FASTQ generation tools

In addition to the direct applications of the simulated VCFs –such as analyzing mutational signatures, benchmarking driver genes detection, and augmenting data to improve existing algorithms– OncoGAN simulations can also be used to generate personalized BAM files for benchmarking genomic analysis tools that operate at the unaligned read level. To illustrate OncoGAN’s compatibility with FASTQ generation tools, we provide a basic script in our GitHub repository that integrates OncoGAN-generated VCF files with a reference genome to produce raw sequencing files using InSilicoSeq (v2.0.1)^39^. This approach can be easily adapted for other FASTQ/BAM generation tools or extended to produce more complex outputs, enabling the creation of an unlimited number of realistic synthetic genomes.

### Open Access Repository of Simulated Cancer Genomes

To provide a foundation for community cancer genome analysis benchmarking efforts, we have used the OncoGAN pipeline to generate 800 simulated cancer genome VCFs and matched CNAs and SVs files across a diverse set of eight tumor types. These simulations are now publicly available for download and redistribution through our project repositories on HuggingFace (https://huggingface.co/datasets/anderdnavarro/OncoGAN-syntheticVCFs) and Zenodo (https://zenodo.org/records/14889626), with additional tumor types and features to be included in future releases as we continue to train and refine the models. Because VCF is a widely accepted standard for mutation calling files and the formats used for CNAs and SVs are the same ones reported in the PCWG study, these datasets can be easily integrated into existing range of bioinformatic tools, facilitating their adoption by researchers in the field.

## Discussion

Here, we present OncoGAN, a generative AI pipeline that leverages deep learning algorithms to simulate highly realistic synthetic cancer genomes. We have demonstrated that OncoGAN reproduces features found in various tumor types with a high level of fidelity while also capturing the significant heterogeneity found across donor populations. By applying the widely-used cancer genome analysis tools SigProfilerExtractor, DeepTumour and ActiveDriverWGS, we showed that the synthetic genomes produced by OncoGAN discovered the same mutational signatures, cell-of-origin signatures, and driver mutation profiles as they do when applied to real data. Finally, we showed that by supplementing real training set data with OncoGAN-simulated data, we could improve the performance of the DeepTumour tumor-type prediction model, suggesting that synthetic genomes could be useful in augmenting the training sets of other genome machine learning applications. In this regard, we successfully worked with tumors that have a small number of donors, such as Lymph-MCLL and Lymph-UCLL, with 35 and 60 donors, respectively. This demonstrates that OncoGAN has the potential to augment datasets for other rare tumor types.

We believe that these highly realistic synthetic cancer genomes can help address some of the current challenges in genomics, such as data accessibility^20–22^, by promoting data sharing while protecting patient confidentiality^40^. We have demonstrated that OncoGAN effectively protects patient privacy by simulating the position and context of mutations independently. Additionally, the training set contains only somatic mutations (with no germline information) from which donor identifiers have been removed. As a result, our synthetic genomes can be made open access and fully available to the research community. This approach aligns with initiatives like the Common Infrastructure for National Cohorts in Europe, Canada, and Africa (CINECA)^41^ or the work by D’Amico et al.^42^, which have synthesized cohort datasets containing phenotypic data. Moreover, data augmentation using synthetic genomes can help resolve class imbalance and small sample size issues^40,43^, providing valuable resources for training new algorithms or improving existing ones, as demonstrated in the DeepTumour example. This approach has also been successfully applied in other fields, including medical image analysis^44^, survival analysis^45,46^ or single-cell data^47,48^.

In recent years, several alternative approaches have been developed for the simulation of synthetic genomes, each employing one of three main strategies. BAMsurgeon^19^, Xome-Blender^49^, and SomatoSim^50^ each use a real sequencing alignment file (BAM) as input, add the alterations, and output the modified BAM along with a list of the incorporated mutations. In contrast, insiM^18^ takes a similar approach by using a BAM file as input, but it outputs paired reads in FASTQ format. Finally, simuG^51^ and Mutation-Simulator^52^ use the reference genome in FASTA format as input, producing an altered FASTA file, which can then be converted into FASTQ reads using any read simulator, such as InSilicoSeq^39^ or Sandy^53^. However, all of these tools primarily focus on integrating random mutations and/or more complex alterations, which do not accurately represent real tumor characteristics. In comparison, OncoGAN is the only method capable of simulating realistic mutational and copy number profiles specific to different tumor types. Moreover, as we have demonstrated, it can be integrated with any of the previous approaches to generate more realistic *in silico* sequencing files, thereby enabling synthetic genomes to address more complex tasks.

Currently, OncoGAN generates synthetic donors for eight tumor types. In the future, we plan to expand this work to include additional tumor types, including those that are less common, extend its capabilities to simulate a broader range of tumor features, such as indel signatures, and also incorporate germline haplotypes. Overall, this will allow us to create realistic simulated raw sequencing data for use in sequence alignment, variant calling, mutational signatures discovery and the detection of copy number alterations and structural variants in cancer genome analysis.

In conclusion, OncoGAN represents a significant advance in the generation of synthetic cancer genomes. It offers a solution to current challenges in data accessibility and privacy while also serving as a powerful tool for enhancing algorithm development and benchmarking.

## Methods

### Data preprocessing

#### Point mutations

The original file used to extract all features for training the different models was the `final_consensus_passonly.snv_mnv_indel.icgc.public.maf.gz` (GRCh37) from the ICGC 25K data portal^4^. This file contains all final mutations detected by whole genome sequencing (WGS) for each donor across the 25 tumor types studied in the PCAWG. Twelve tumor types were discarded due to an insufficient number of samples for model training (<50 donors). From the remaining thirteen tumor types, we selected eight, ensuring a heterogeneous dataset based on factors such as the tumor mutational burden, the number of available donors and the tumor subtype. The selected studies were: Breast-AdenoCa, CNS-PiloAstro, Eso-AdenoCa, Kidney-RCC, Liver-HCC, Lymph-CLL, Panc-Endocrine and Prost-AdenoCa. For each mutation, we retained the following information: study (tumor type), donor ID, chromosome, position, length, type of mutation, reference and alternative bases, trinucleotide context and variant allele frequency. Mutations were categorized into five groups: single nucleotide polymorphisms (SNPs; A > C), double nucleotide polymorphisms (DNPs; AA > CC), triple nucleotide polymorphisms (TNPs; AAA > CCC), insertions (INS; A > AA) and deletions (DEL; AA > A). Mutations without variant allele frequency were excluded.

To associate each SNP with its corresponding mutational signature (single base substitution; SBS) from the COSMIC catalogue (https://cancer.sanger.ac.uk/signatures/sbs/), individual variant call format files were extracted for each donor. SigProfilerExtractor^34^ (v1.1.21) was then run, individually for each tumor type, using its default configuration except for the `--maximum_signatures` option, which was set to 8. The `De_Novo_MutationType_Probabilities.txt` file was used to sample the most probable signature each SNP belongs to, based on donor ID and mutational context. SBSs present in 5% or fewer (rounded) of donors were removed from the dataset, as more data would be required to accurately train the models.

Since categorical columns in tabular data are converted into one-hot encoding, we simplified the dataset by removing the chromosome column. Thus, positions were transformed into a continuous range along the genome (e.g., if chromosome 1’s length is 1000, position 1 on chromosome 2 would be 1001).

The preprocessed file was then used to prepare the training files for all the models described below except for CNA and SV related models.

#### Copy number alterations and structural variants

The training files for CNA and SV simulations were created using the original datasets available in the `consensus.20170119.somatic.cna.icgc.public` and `final_consensus_sv_bedpe_passonly.icgc.public` folders, respectively, from the ICGC 25K data portal^4^. Due to the limited number of events, all tumor types with more than 30 donors were included to ensure sufficient data for training the models. While all donors were used, the tumor type of origin was retained as a feature in the training file for CNA-related events, enabling tumor-specific alteration simulations. However, for SV-related events, this feature was excluded due to the insufficient number of alterations within each tumor type. The lengths of the events were calculated using their start and end positions and then transformed using natural logarithms to reduce variability and improve data distribution. These processed datasets were used to generate four distinct training files, which are described below.

### Model training

Detailed hyperparameters settings for all the models can be found in Supplementary Table S12.

#### Donor characteristics

Donor characteristics for point mutations refer to the number and types of mutations and signatures present in each donor. To generate this dataset, we counted the number of each mutation type per donor, replacing SNPs with the assigned SBSs to avoid duplicating information in the training file. We trained a separate model for each of the tumor types studied, using CTAB-GAN+^29^ with 720 different hyperparameter combinations (epochs: 100 to 500 in increments of 20; batch size: 10 to 30 in increments of 5; learning rate: 0.001 to 0.01 in increments of 0.001). The best model or models were selected based on the similarity of scatter and density plots comparing the simulations to the original data. Given the complexity of tumors –where different mutational types and signatures interact in varied ways– and the limited data available for certain tumor types, we refined the simulations by combining models trained with different hyperparameter settings and applying tumor-specific ad-hoc filters. This approach allowed us to better capture low-frequency mutational signature patterns, ensuring that the generated data accurately resembled the original distributions.

For copy number alterations and structural variants, donor characteristics include features such as the tumor type, the number of CNAs and each type of SV (deletions, duplications, inversions, and translocations), the total length of aberrant CNA segments, and the total number of mutations per donor. The total number of mutations is used to guide the simulation of CNAs and SVs, ensuring they match the donor’s mutational profile. Given the simplicity of the data, we trained a single CTAB-GAN+ model to simulate donor characteristics for CNAs and SVs, then the tumor type feature can be used to specifically simulate samples for the desired tumor.

#### Driver genes

Since OncoGAN randomly generates positions and trinucleotide contexts, the resulting mutations represent only passenger mutations. Therefore, OncoGAN cannot directly simulate driver mutations, as it lacks the biological information necessary for this task. To address this limitation, we first selected a list of driver genes published by PCAWG and available at the ICGC 25K data portal (`TableS1_compendium_mutational_drivers.xlsx`). We classified mutations in driver genes as either “coding” or “non-coding,” creating a new ID (e.g., TP53_coding / TP53_intron). Driver IDs were filtered to include only those present in at least 5% of donors for coding mutations and 10% for non-coding mutations. The remaining mutations were used to create a database from which mutations could be drawn during the simulation.

However, driver IDs cannot be randomly selected, as some drivers are more common than others, and some tend to occur together or be mutually exclusive. To simulate these driver’s intercorrelation, we trained a specific model. The training file for this model consisted of a count of how many driver mutations each donor had for each driver ID.

The same approach used for training donor characteristics models was applied here, with one model trained for each tumor type (CTAB-GAN+^29^ with 720 hyperparameter combinations). The best models were manually selected based on the frequency of mutations for each driver ID and the correlation between driver’s intercorrelations.

#### Variant allele frequency

Variant allele frequency was simulated using a straightforward approach since it is related to tumor purity and independent of the donor and mutation characteristics described above. First, we calculated the mean VAF for each donor and ranked them into 100 VAF bins (1% increments, from 0 to 1) to determine the percentage of donors that fall into each bin, independently for each tumor type. Then, within each of these ranks, we selected the mutations belonging to donors within each bin and further ranked them according to 40 VAF bins (2.5% VAF increments, from 0 to 1), to calculate the percentage of mutations in each bin. To simulate a donor, we first sample a mean donor VAF from the first file using the calculated probabilities and then sample mutation VAFs using the probabilities calculated in the second file for that specific mean donor VAF.

#### Genomic position

The genomic position of mutations is a variable with a very wide range, spanning from 1 to 3 billion base pairs (3Gbp), making it difficult for generative models to capture this variability. To address this, we performed a two-step discretization of the genome. First, we compressed the 3Gbp genome into 30Mbp (a 1:100 ratio) to reduce the complexity, as single-base resolution is not necessary for passenger mutations. Next, we divided the compressed genome into 105 bins of 0.3Mbp each, matching each mutation’s position to its corresponding bin, discarding bins with fewer than two mutations. We then used the CTGAN^27^ model to simulate new bins at the same frequency found in the original data. In the second step, we trained a small tabular variational autoencoder (TVAE)^27^ model independently for each bin, using the positions rather than the bins as input for the training. Finally, the simulated positions were expanded to the actual genome size. The use of both architectures is necessary because the GAN model better captures the overall frequency of the bins, but it is unable to accurately reproduce the specific mutation positions when working with a larger range.

The same strategy was used to train a model to simulate the position of the structural variants.

#### Mutational signatures

To simulate the mutations, we prepared a training file containing the following features: position, length, VAF, reference and alternative contexts, and signature. The context is represented with individual bases in separate columns (r.ctx1, r.ctx2, r.ctx3, a.ctx1, a.ctx2, a.ctx3). Although we trained specific models for genomic position and VAF, we included these features in this training file as well, as they stabilized and improved model performance, preventing exploding issues in the gradient penalty during the GAN model’s training phase.

We used a CTAB-GAN+^29^ architecture for these models, defining the variables as follows: ‘r.ctx1’, ‘r.ctx2’, ‘r.ctx3’, ‘a.ctx1’, ‘a.ctx2’, ‘a.ctx3’, and ‘signature’ as categorical; length as a mixed type (with 0 representing “no indel” as a categorical value, >0 representing insertion size and <0 for deletion size); position as an integer; and VAF as a default float. Hyperparameters were selected based on tumor characteristics, with individual optimization for each tumor type. Batch size was set to the mean number of mutations per donor, the learning rate to 2e-4, and epochs ranged from 280 to 3200 to adapt the training time to an average of five days. The ‘signature’ variable was not simulated directly by the GAN model but was instead predicted by the auxiliary component of the CTAB-GAN+ model, based on the other simulated variables. This classifier was trained using a test ratio of 0.3. Since the goal of these models is to reproduce mutational signatures, the model’s performance was tested by simulating donors and running SigProfilerExtractor (v1.1.21) to detect the simulated signatures. During the training phase, we experimented with different configurations and observed that the use of fewer epochs caused SigProfilerExtractor to detect new signatures (artifacts) not present in the catalog.

#### Copy number alterations and structural variants

Two independent models were trained using a CTAB-GAN+ architecture to simulate copy number alterations and structural variants. For CNAs, the training file includes the total length of altered segments (log scale), the copy number state of both alleles, and the tumor type to which the alteration belongs. Events with rare copy number states (< 0.5%) were excluded. The SV training file includes the genomic position where the event starts, the event length (log scale), strand information, and the SV class. For translocations, the length represents the linear distance between the two breakpoints of the chromosomes involved in the event, allowing the model to reproduce interactions between close or distant chromosomes with a frequency similar to that observed in the real data. However, the original PCAWG files did not provide information about the actual sizes of the two fragments involved in translocations.

### Quality control metrics

#### Donor characteristics

To assess the similarity between real and simulated populations, we calculated the histogram intersection distance for each type of mutation using the HistogramTools (v0.3.2) R package. First, we determined the number of each mutation type per donor. Then, we generated histograms for both the real and simulated datasets using the same bins (nbin = 10, spanning from the minimum to maximum values for each mutation type). Histogram counts were normalized by the total number of donors to make the histograms comparable. We calculated three different intersection distances to evaluate the differences between the populations: i) The histogram intersection distance between the PCAWG and OncoGAN datasets. ii) A bootstrap analysis (n = 1,000) for the above comparison to calculate confidence intervals. iii) A control analysis in which the PCAWG dataset was split into two random subgroups, and the intersection distance was calculated within them (n = 1,000). The closer the intersection distance is to 0, the more similar the two populations are.

To compare the simulations not only to their corresponding real tumors but also to other tumor types, we calculated the histogram intersection distances across the eight studied tumor types. In this case, we used the total number of mutations per donor as the comparison metric. Using the minimum and maximum values across the eight tumor types, we defined 100 bin ranges to ensure consistency across all comparisons. We compared the PCAWG dataset against itself to determine the true relationship between the tumor types and then compared the OncoGAN dataset against PCAWG. For a more realistic comparison when calculating the intersection distance within the same tumor type for the PCAWG dataset (e.g., Breast-AdenoCa vs. Breast-AdenoCa), we randomly split the dataset into two subgroups and performed the intersection distance calculation 200 times, reporting the mean intersection distance. Finally, to evaluate the similarity between real and simulated distributions, we calculated the Pearson correlation.

#### Driver genes

To verify the accuracy of simulating driver gene co-occurrence, we calculated the percentage of donors harboring each possible 1-vs.-1 driver combination for both the real and simulated datasets. The four possible combinations were: i) neither driver was present; ii) only “driver A” was present;

iii) only “driver B” was present; and iv) both drivers were present. Pearson correlation was then calculated independently for each of the eight studied tumor types, considering all combinations. The number of selected drivers and combinations can be found in Supplementary Table S4.

To further assess whether the introduced mutations are identified as driver genes at the population level, as observed in real data, we used the software ActiveDriverWGS (v1.2.1)^36^. The analysis was performed independently for both the PCAWG and OncoGAN datasets, using ActiveDriverWGS’s default cancer gene list and configuration. Genes were classified as drivers if their FDR-corrected p-value was below 0.05.

#### Variant allele frequency

To confirm that the mean variant allele frequency per donor was simulated according to the real distribution, we performed a two-sided Wilcoxon test comparing real and simulated donor VAFs independently for each tumor type. Additionally, a two-sided F-test was conducted to compare the variances of real and simulated populations.

#### Genomic position

To evaluate how accurately the locations of mutations were simulated according to patterns found in real tumors, we employed three approaches: i) The human genome was divided into 3,097 bins (1Mbp) and the percentage of mutations in each bin for both the real and simulated datasets was calculated. Pearson correlation was then estimated for each of the studied tumor types using these values. ii) A linear regression was performed to compare the PCAWG dataset against itself, and OncoGAN vs. PCAWG datasets to determine whether there were differences in the simulation of mutation positions between regions with lower or higher density of mutations, reporting the R^2^ value. When comparing the same tumor type (e.g. Breast-AdenoCa vs. Breast-AdenoCa), we split the PCAWG dataset into two random subgroups and ran the linear regression 200 times, reporting the mean values. iii) t-SNE visualization (Rtsne package v0.17 with default parameters) was used to cluster real and synthetic donors based on the genomic distribution of their mutations (absolute values for 1Mbp windows).

#### Mutational signatures

To detect mutational signatures present in OncoGAN simulations, we employed the same preprocessing approach used for the PCAWG dataset, using SigProfilerExtractor (v1.1.21) with its default configuration, except the `--maximum_signatures` option was set to 8. Signatures were then manually compared to determine whether all expected signatures were simulated and to check for any new signatures or artifacts. We also compared the percentage of donors harboring the mutations with the corresponding percentages found in the real data. To investigate discrepancies between OncoGAN simulations and SigProfiler detections, we visualized the percentage of mutations –both simulated and detected– associated with each signature for each donor. Additionally, we examined discrepancies in the primary trinucleotide contexts (≥0.5%) present in each signature by comparing COSMIC (v3.4) (https://cancer.sanger.ac.uk/signatures/downloads/) with their usage in the PCAWG training file and the dataset generated by OncoGAN. Pearson correlations were calculated to assess the similarity between the patterns.

#### Indel distribution

To evaluate whether the distribution of indel sizes in the simulations closely matched that observed in the real data, we performed a Wilcoxon test between the two datasets for each indel size. For this analysis, we calculated the percentage that each indel length contributed to the total number of indels for each donor. The analysis was restricted to indels up to 5 nucleotides in length, as these account for an average of 88.5% of all indels.

#### Mutational consequences

The Docker version of the Ensembl Variant Effect Predictor tool (v112)^35^ was used to predict the effects of real and simulated mutations. The cache for the *Homo sapiens* GRCh37 version of the genome was downloaded from the Ensembl SFTP server (https://ftp.ensembl.org/pub/release-112/variation/indexed_vep_cache/), and results were obtained only for canonical transcripts using the following command: `vep --offline --format vcf --dir_cache /cache/ -- force_overwrite --total_length --numbers --ccds --canonical --biotype --pick - -no_stats --assembly GRCh37.` To enhance plot readability, Y axes were broken using the R package ggbreak (v0.1.2)^54^.

#### Copy number alterations and structural variants

The similarity between real and simulated CNA and SV profiles was compared using four chromosomal instability metrics adapted from the CINmetrics package^38^. For copy number alterations, we used the copy number abnormality score^55^, which is defined as the number of segments with an altered copy number (i.e., where the major and minor alleles differ from one). Additional metrics include the mean length per altered segment, the fraction of the genome altered (calculated as the total length of altered segments normalized by genome size), and the total aberration index (TAI)^56^, which is calculated as:

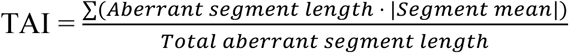

Where segment mean is defined as:

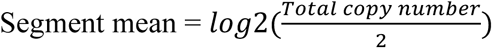

For structural variants, two metrics were used: the number of SVs and the mean length of the SV events per donor. In all cases, the Wilcoxon test was used to compare the means of the different metrics between the real and simulated donors.

### DeepTumour testing and training

DeepTumour is a fully connected feedforward deep neural network that in its latest version classifies 29 common cancer types based on mutation positional and context distributions (https://deeptumour.oicr.on.ca/)^37^. To generate the positional distributions, DeepTumour divides the genome into 2,897 bins (1Mbp) and counts the number of somatic mutations within each bin. Additionally, it calculates the distribution of 150 different mutation types in their single (e.g. C > G), dinucleotide (e.g. AC > AG), and trinucleotide contexts (e.g. ACA > AGA). The count for each mutation type is then normalized by the total number of SNVs in the sample and then transformed into z-scores for each feature. These positional and contextual mutation distributions result in a total of 3,047 features. We used this version of DeepTumour as a quality control tool to predict the tumor type of origin for our simulations, providing the predicted tumor type and the probability of the sample belonging to any other possible tumor types. To explain the Eso-AdenoCa results, we performed a PCA using the PCAtools (v2.14.0) R package and the main donor characteristics (number of mutations for each type and mutational signature). For Lymph-MCLL, we compared their genomic position metrics with those from Lymph-BNHL.

To observe changes in DeepTumour’s performance with simulated samples generated by OncoGAN, we trained a new model using the same architecture as for the baseline model. Originally for the baseline model, the 2016 release of 2,778 PCAWG data was filtered for tumor types with 18 or more samples, resulting in 29 cancer types. These samples were then split into 80% training data, 10% validation data, and 10% testing data, repeated five times since the model was trained following a 5-fold cross validation. Using each of the five training partitions, the model was trained with Adam50, employing a batch size of 32 for 400 epochs in PyTorch 1.9.1 with CUDA 10.2 support. Hyperparameter optimization was conducted by improving the model’s performance on the corresponding split of validation data using the ‘gp_minimize’ function from the scikit-optimize 0.9.0 Python library. The new model was trained using a consistent approach, incorporating 100 additional simulated samples generated by OncoGAN for each tumor type studied in the original 80% of the training data. The code was written in Python 3.7.16.

To compare the performance of DeepTumour trained with simulated samples to the baseline model, we evaluated traditional performance metrics, including accuracy, precision, recall, and F1 score, independently using the corresponding test partitions. To calculate the overall accuracy, we calculated the proportion of correct classification. This metric is applied only when summarizing the performance of the classifier across all 29 tumor types. Recall was calculated as the proportion of samples from a specific cancer type that were correctly classified as that type, while precision measures the proportion of samples assigned to a particular type that truly belong to that type. The F1 score represents the harmonic mean of recall and precision. When reporting independent Lymph-MCLL and Lymph-UCLL metrics, Lymph-CLL donors and misclassified donors from other tumor types were divided based on the presence or absence of the SBS9 mutational signature, respectively.

### Availability of data and materials

The OncoGAN software is publicly available on GitHub (https://github.com/LincolnSteinLab/oncoGAN) and DockerHub (https://hub.docker.com/r/oicr/oncogan) to enhance reproducibility. The training files, trained models, and the 800 simulated donors used in this paper have been uploaded to HuggingFace (https://huggingface.co/collections/anderdnavarro/oncogan-67110940dcbafe5f1aa2d524) and Zenodo (https://zenodo.org/records/14889626).

## Acknowledgements

We thank Matus Medo, Quaid Morris, Valli Subasri and Phil Fradkin for their useful suggestions. This work was performed using High Performance Computing resources from the Ontario Institute for Cancer Research.

## Contribution

D-N.A.: Conceptualization (equal); Data curation (lead); Formal analysis (lead); Investigation (lead); Methodology (lead); Project administration (equal); Software (lead); Visualization (lead); Writing – Original Draft Preparation (lead); Writing – Review & Editing (equal). Z.X.: Formal analysis (supporting); Methodology (supporting); Visualization (supporting); Writing – Original Draft Preparation (supporting); Writing – Review & Editing (supporting). J.W.: Formal analysis (supporting); Methodology (supporting). W.B.: Funding acquisition (supporting); Project administration (supporting); Supervision (supporting); Writing – Review & Editing (supporting). S.L.: Conceptualization (equal); Funding acquisition (lead); Project administration (equal); Resources (lead); Supervision (lead); Writing – Review & Editing (equal)

## Supplementary Data

**Supplementary Figure S1.** Scatter and density plots comparing the number of specific mutation types to the total number of mutations for each donor, with real donors from PCAWG shown in orange, and simulated donors from OncoGAN shown in black. In all cases, the distributions are very similar, although there are minor challenges in simulating donors with very low mutational burdens in Eso-AdenoCa and Kidney-RCC tumor types.

**Supplementary Figure S2.** Violin plots illustrating the distribution of intersection distances between two randomly sampled populations from the PCAWG dataset (1000 interations) and the scores comparing OncoGAN simulations to the actual dataset. For most mutation and tumor types, the simulations correspond closely with the obtained results from comparisons between two subpopulations in the real data, indicating a high degree of similarity between the PCAWG and OncoGAN distributions. Lower scores indicate greater similarity between the populations.

**Supplementary Figure S3.** Comparative density and box plots showing donor’s distributions of mean variant allele frequency (VAF) across eight tumor types. Each panel compares the VAFs from real samples (PCAWG) in orange and simulated samples (OncoGAN) in black. Notably, only the CNS-PiloAstro panel shows a statistically significant difference, which may not be biologically relevant as both distributions are very similar with median VAFs of 0.222 and 0.238, respectively. This underscores the accuracy of the OncoGAN simulations in reflecting real data. The Wilcoxon test was used to compare the groups. *NS.: p-value > 0.05; *: p-value <= 0.05; **: p-value <= 0.01; ***: p-value <= 0.001*.

**Supplementary Figure S4.** Density plots for the variant allele frequency (VAF) for each type of mutation and tumor. The densities are very similar between real (orange) and simulated (black) VAFs. The number of each type of mutation in each dataset is also reported. The greatest differences are observed in TNPs due to its low frequency.

**Supplementary Figure S5.** Driver correlation analysis plot for the seven tumor types. The X-axis shows all possible combinations between two driver genes, with each dot representing one 1-vs-1 combination. The Y-axis represents the percentage of donors in which that combination occurs. The number of drivers used and the total number of combinations are reported in the caption for each tumor type. The Pearson coefficient, with values exceeding 0.9 in almost all tumors, indicates a strong simulation of driver relationships. The data used to create this plot, including all possible driver combinations and their values, is listed in SuppTableS3.

**Supplementary Figure S6.** Histograms displaying the total percentage of mutations across the genome in 1Mbp bins for the remaining seven tumor types. Real donors are shown in orange, and simulated ones in black, with differences highlighted in green. Pearson correlations between PCAWG and OncoGAN genomic profiles are provided for each specific plot.

**Supplementary Figure S7.** Scatter plot comparing the percentage of mutations in each 1Mbp genomic region between two samples from the PCAWG dataset (left) and OncoGAN simulations against the entire PCAWG dataset (right) for each of the remaining tumor types. Each dot represents a region, with color indicating density; lighter colors mean higher densities. R^2^ values are displayed for each of the comparisons.

**Supplementary Figure S8.** Scatter and density plots comparing the number of specific signature mutations to the total number of mutations for each donor, with real donors from PCAWG shown in orange, and simulated donors from OncoGAN shown in black. OncoGAN values correspond to those directly simulated by the tool.

**Supplementary Figure S9.** Mutational signatures distributions detected by SigProfiler showing the number of somatic mutations per megabase and the percentage of donors exhibiting each signature for the remaining tumor types. Dots represent individual donors; the red line illustrates the mean number of somatic mutations per megabase. *SBS, single base substitutions*.

**Supplementary Figure S10.** Comparison of the mutational pattern for the SBS30 signature in the Eso-AdenoCa tumor type across the PCAWG, OncoGAN, and COSMIC datasets. A) For all trinucleotide contexts the percentage each specific context contributes to the signature. B) Focusing on the C>T context, as it is the predominant altered context in this signature.

**Supplementary Figure S11.** Comparison of the percentage of mutations directly simulated (OncoGAN, green) and detected (SigProfiler, blue) for each mutational signature across donors within the eight studied tumor types. Red dashed line indicates the lowest percentage of mutations detected by SigProfiler. *SNP, Single nucleotide polymorphism*.

**Supplementary Figure S12.** Boxplots comparing indel length distribution between PCAWG (orange) and OncoGAN (black). Negative X values represent deletions, and positive X values represent insertions. The Y-axis shows the contribution (%) of each specific indel length relative to the total number of indels per donor. Only indels up to a size of 5 are plotted. The Wilcoxon test was used to compare the groups. *NS.: p-value > 0.05; *: p-value <= 0.05; **: p-value <= 0.01; ***: p-value <= 0.001*.

**Supplementary Figure S13.** Boxplots comparing the frequency of predicted effects for real (orange) and simulated (black) mutations per donor using the Variant Effect Predictor (VEP) tool. The X-axis lists the possible mutation consequences considered by VEP, while the Y-axis shows the percentage of mutations corresponding to each consequence per donor.

**Supplementary Figure S14.** Comparison of real Lymph-MCLL and Lymph-BNHL tumor types illustrating their similarity in terms of genomic mutation profiles. A) Histogram displaying the total percentage of mutations across the genome in 1Mbp bins. Real donors are shown in orange, and simulated ones in black, with differences highlighted in green. The Pearson correlation is 0.941, indicating a high degree of similarity. B) Scatter plot comparing the percentage of mutations in each 1Mbp genomic region between both tumor types. Each dot represents a region, with color indicating density; lighter colors indicate higher densities.

**Supplementary Figure S15.** DeepTumour heatmap displaying the accuracy of the baseline and the new classifier using a held-out portion of the PCAWG data set for evaluation (5-fold cross-validation). Baseline model was trained using only PCAWG samples, whereas the new model employs a mix of real and synthetic samples. Each row corresponds to the true tumor type and columns correspond to the class predictions emitted by DeepTumour. Cells are labeled with the percentage of donors of a particular type that were classified by DeepTumour as a particular type. The recall and precision of each classifier are shown in the color bars at the top and left sides of the matrix. All values represent the mean of 5 runs using selected data set partitions.

**Supplementary Figure S16.** Scatter and density plots showing the relationship between the total number of mutations per donor and two chromosomal instability scores: the total number of aberrant segments and their mean length. As shown, the distributions of real (orange) and simulated (black) donors are highly similar.

**Supplementary Figure S17.** Chromosomal instability scores measuring the similarity of inversion and translocation SVs between real (orange) and simulated (black) donors. A) Number of h2hINV, t2tINV, and TRA events per donor. B) Mean length of SVs in base pairs. Error bars represent the standard deviation of the dataset. The Wilcoxon test was used to compare the groups. *NS.: p-value > 0.05; *: p-value <= 0.05; **: p-value <= 0.01; ***: p-value <= 0.001. h2hINV, head-to-head inversion; t2tINV, tail-to-tail inversion; TRA, Translocation*.

**Supplementary Table S1**. Percentage of duplicated mutations within PCAWG and OncoGAN datasets, as well as between them for each tumor type.

**Supplementary Table S2**. Mean intersection distances and confidence intervals for the distribution of number of mutations, comparing PCAWG (Sampled population 1 vs. Sampled population 2) with OncoGAN (OncoGAN vs. PCAWG) donors.

**Supplementary Table S3.** Results of Wilcoxon and F-tests comparing the distributions of donor variant allele frequencies and the equality of their variances (PCAWG vs. OncoGAN).

**Supplementary Table S4.** Driver correlation analysis table displaying the percentage of donors that carry each combination of driver mutations.

**Supplementary Table S5.** Linear regression results showing the R² for comparisons between two samples from the PCAWG dataset (n=200) and OncoGAN simulations versus the PCAWG regarding the percentage of mutations distributed across the genome (1Mbp bin size).

**Supplementary Table S6.** Pearson coefficients and confidence intervals to measure the similarity of context contribution for mutational signatures between OncoGAN and PCAWG datasets and the COSMIC ground truth. Only contexts whose contribution is equal to or greater than 0.5% of the signature are used.

**Supplementary Table S7.** For each cancer type and mutational signature, the number of donors in which the specific signature was simulated by OncoGAN and the number of donors in which the signature was detected by SigProfiler. The last two columns present, for each specific signature, the mean total number of simulated mutations for donors where the signature was also detected by SigProfiler and for donors where the signature was not detected.

**Supplementary Table S8.** Pearson coefficients representing the correlation between the percentages of each mutational signature simulated (OncoGAN) and those detected (SigProfiler) in simulated donors. study_error indicates the discrepancy between the number of donors that were intended to be simulated with a specific mutational signature and those actually detected.

**Supplementary Table S9.** Performance of ActiveDriverWGS for the discovery of cancer driver genes in both PCAWG and OncoGAN datasets. Genes without p-value are genes not present in the default list of cancer genes used by ActiveDriverWGS.

**Supplementary Table S10.** DeepTumour baseline model predictions for the OncoGAN dataset and 5-fold cross-validation results for the PCAWG dataset.

**Supplementary Table S11.** Chromosomal instability metrics for different tumor types in the PCAWG and OncoGAN datasets. For each tumor type, results are shown separately for copy number alterations (CNA) and structural variants (SV). Metrics include the number of segments, segment size (Mb), fraction of the genome altered, and total aberration index. Values represent the mean, with standard deviation in parentheses.

**Supplementary Table S12.** List of hyperparameters used to train the models.

