## Supplemental Figures for "*In silico* generation of synthetic cancer genomes using generative AI"

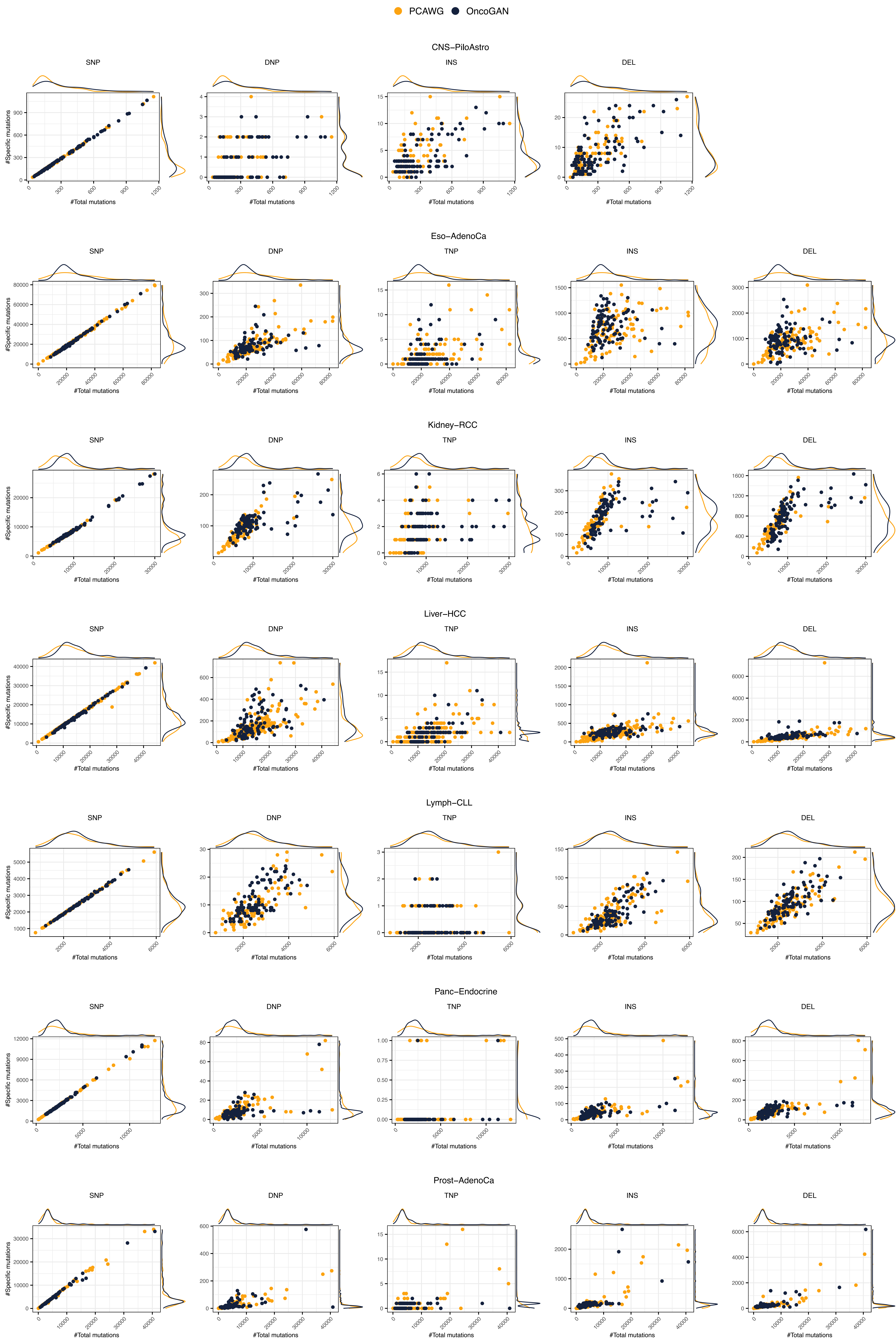
PCAWG
OncoGAN
CNS−PiloAstro
SNP
DNP
INS
DEL
Eso−AdenoCa
TNP
Kidney−RCC
Liver−HCC
Lymph−CLL
Panc−Endocrine
Prost−AdenoCa
#Specific mutations
#Total mutations

**Supplementary Figure S1.** Scatter and density plots comparing the number of specific mutation types to the total number of mutations for each donor, with real donors from PCAWG shown in orange, and simulated donors from OncoGAN shown in black. In all cases, the distributions are very similar, although there are minor challenges in simulating donors with very low mutational burdens in Eso-AdenoCa and Kidney-RCC tumor types.

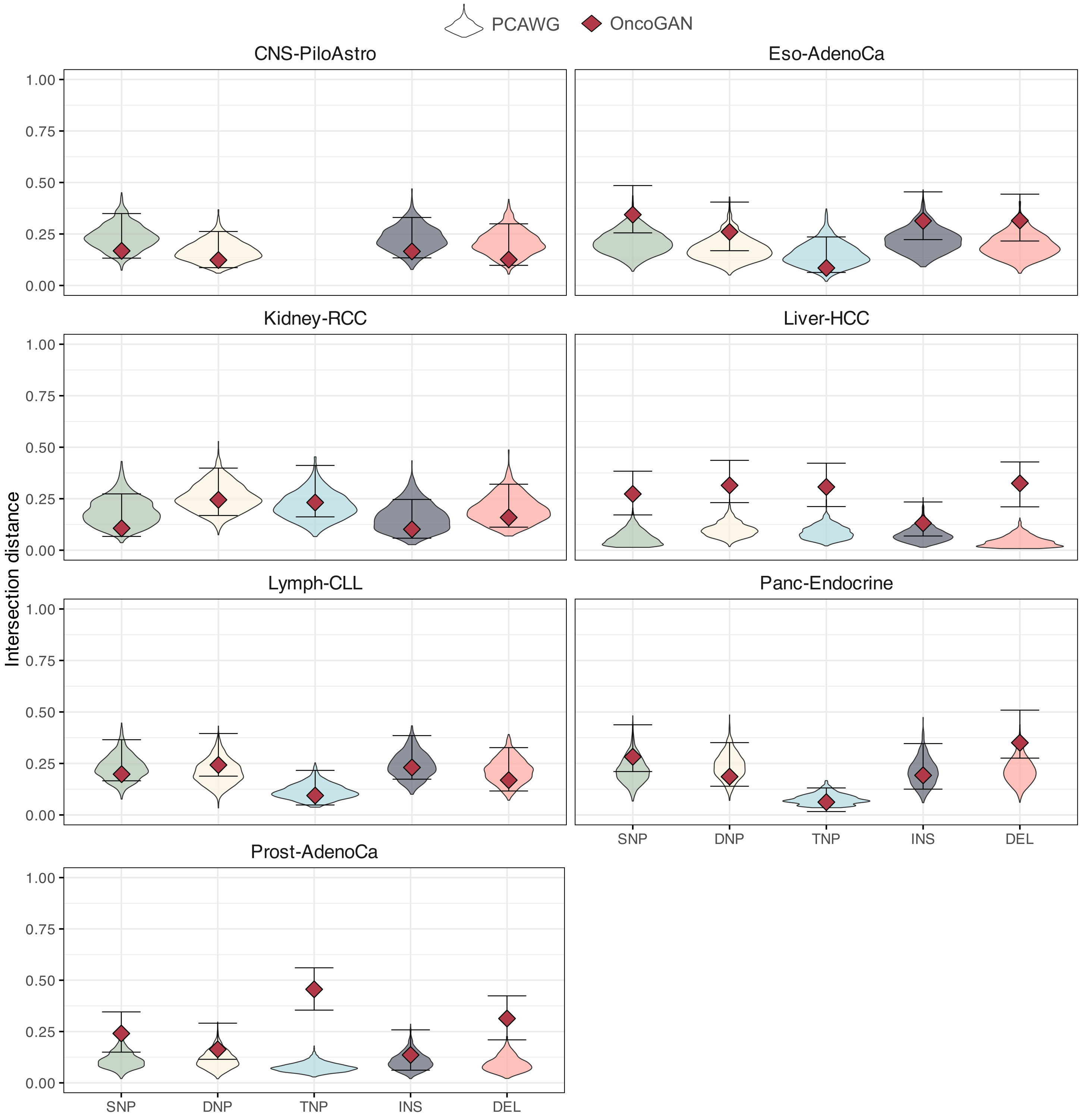

PCAWG
OncoGAN
CNS-PiloAstro
Eso-AdenoCa
Kidney-RCC
Liver-HCC
Lymph-CLL
Panc-Endocrine
Prost-AdenoCa
Intersection distance
1.00
0.75
0.50
0.25
0.00
SNP
DNP
TNP
INS
DEL

**Supplementary Figure S2.** Violin plots illustrating the distribution of intersection distances between two randomly sampled populations from the PCAWG dataset (1000 interations) and the scores comparing OncoGAN simulations to the actual dataset. For most mutation and tumor types, the simulations correspond closely with the obtained results from comparisons between two subpopulations in the real data, indicating a high degree of similarity between the PCAWG and OncoGAN distributions. Lower scores indicate greater similarity between the populations.

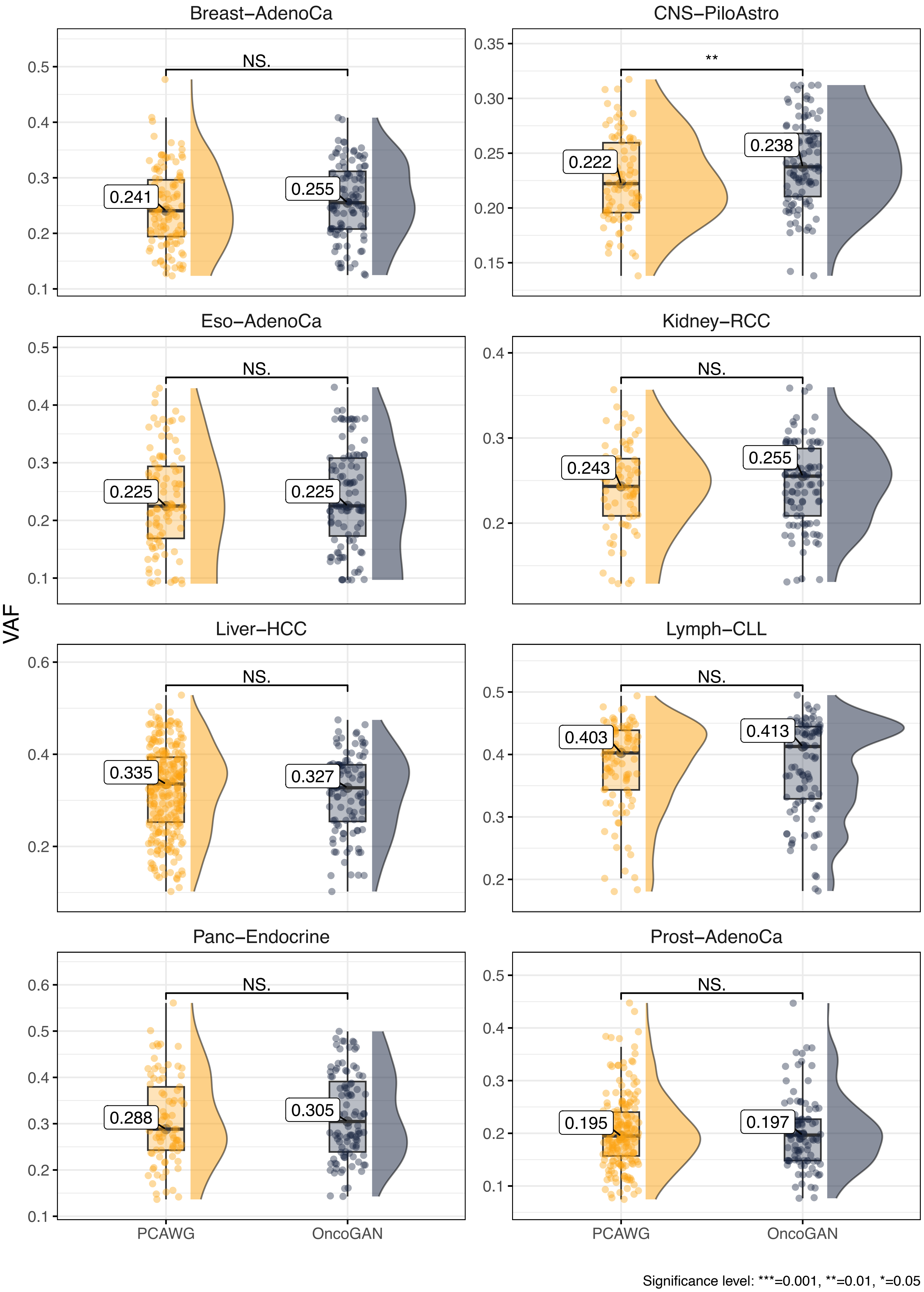
Breast−AdenoCa
NS.
0.241
0.255
CNS−PiloAstro
**
0.222
0.238
Eso−AdenoCa
NS.
0.225
0.225
Kidney−RCC
NS.
0.243
0.255
Liver−HCC
NS.
0.335
0.327
Lymph−CLL
NS.
0.403
0.413
Panc−Endocrine
NS.
0.288
0.305
Prost−AdenoCa
NS.
0.195
0.197
VAF
PCAWG
OncoGAN
PCAWG
OncoGAN
Significance level: ***=0.001, **=0.01, *=0.05

**Supplementary Figure S3.** Comparative density and box plots showing donor's distributions of mean variant allele frequency (VAF) across eight tumor types. Each panel compares the VAFs from real samples (PCAWG) in orange and simulated samples (OncoGAN) in black. Notably, only the CNS-PiloAstro panel shows a statistically significant difference, which may not be biologically relevant as both distributions are very similar with median VAFs of 0.222 and 0.238, respectively. This underscores the accuracy of the OncoGAN simulations in reflecting real data. The Wilcoxon test was used to compare the groups. *NS.: p-value > 0.05; *: p-value <= 0.05; **: p-value <= 0.01; ***: p-value <= 0.001.*

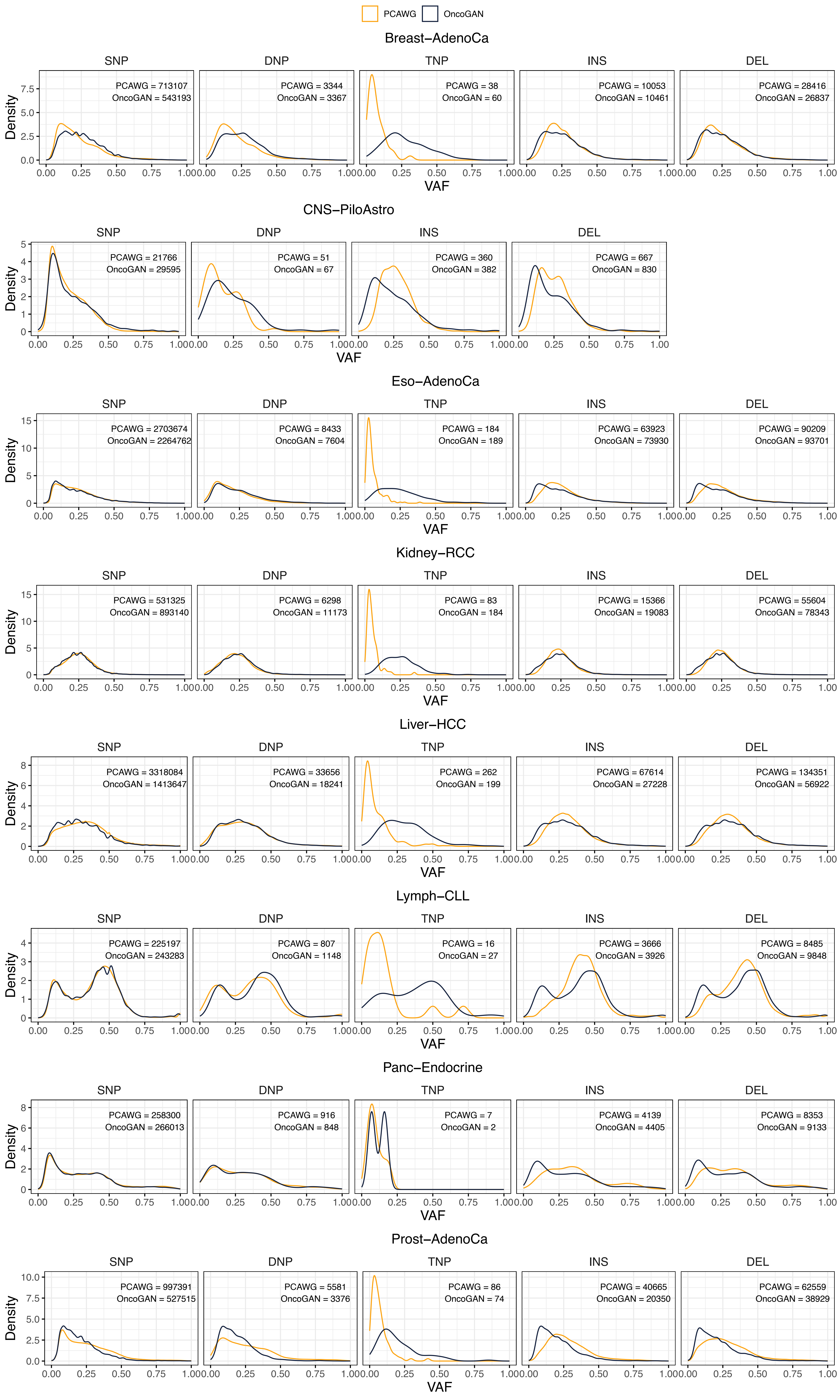
PCAWG
OncoGAN
Breast−AdenoCa
SNP
PCAWG = 713107
OncoGAN = 543193
DNP
PCAWG = 3344
OncoGAN = 3367
TNP
PCAWG = 38
OncoGAN = 60
INS
PCAWG = 10053
OncoGAN = 10461
DEL
PCAWG = 28416
OncoGAN = 26837
CNS−PiloAstro
SNP
PCAWG = 21766
OncoGAN = 29595
DNP
PCAWG = 51
OncoGAN = 67
INS
PCAWG = 360
OncoGAN = 382
DEL
PCAWG = 667
OncoGAN = 830
Eso−AdenoCa
SNP
PCAWG = 2703674
OncoGAN = 2264762
DNP
PCAWG = 8433
OncoGAN = 7604
TNP
PCAWG = 184
OncoGAN = 189
INS
PCAWG = 63923
OncoGAN = 73930
DEL
PCAWG = 90209
OncoGAN = 93701
Kidney−RCC
SNP
PCAWG = 531325
OncoGAN = 893140
DNP
PCAWG = 6298
OncoGAN = 11173
TNP
PCAWG = 83
OncoGAN = 184
INS
PCAWG = 15366
OncoGAN = 19083
DEL
PCAWG = 55604
OncoGAN = 78343
Liver−HCC
SNP
PCAWG = 3318084
OncoGAN = 1413647
DNP
PCAWG = 33656
OncoGAN = 18241
TNP
PCAWG = 262
OncoGAN = 199
INS
PCAWG = 67614
OncoGAN = 27228
DEL
PCAWG = 134351
OncoGAN = 56922
Lymph−CLL
SNP
PCAWG = 225197
OncoGAN = 243283
DNP
PCAWG = 807
OncoGAN = 1148
TNP
PCAWG = 16
OncoGAN = 27
INS
PCAWG = 3666
OncoGAN = 3926
DEL
PCAWG = 8485
OncoGAN = 9848
Panc−Endocrine
SNP
PCAWG = 258300
OncoGAN = 266013
DNP
PCAWG = 916
OncoGAN = 848
TNP
PCAWG = 7
OncoGAN = 2
INS
PCAWG = 4139
OncoGAN = 4405
DEL
PCAWG = 8353
OncoGAN = 9133
Prost−AdenoCa
SNP
PCAWG = 997391
OncoGAN = 527515
DNP
PCAWG = 5581
OncoGAN = 3376
TNP
PCAWG = 86
OncoGAN = 74
INS
PCAWG = 40665
OncoGAN = 20350
DEL
PCAWG = 62559
OncoGAN = 38929
Density
VAF

**Supplementary Figure S4.** Density plots for the variant allele frequency (VAF) for each type of mutation and tumor. The densities are very similar between real (orange) and simulated (black) VAFs. The number of each type of mutation in each dataset is also reported. The greatest differences are observed in TNPs due to its low frequency.

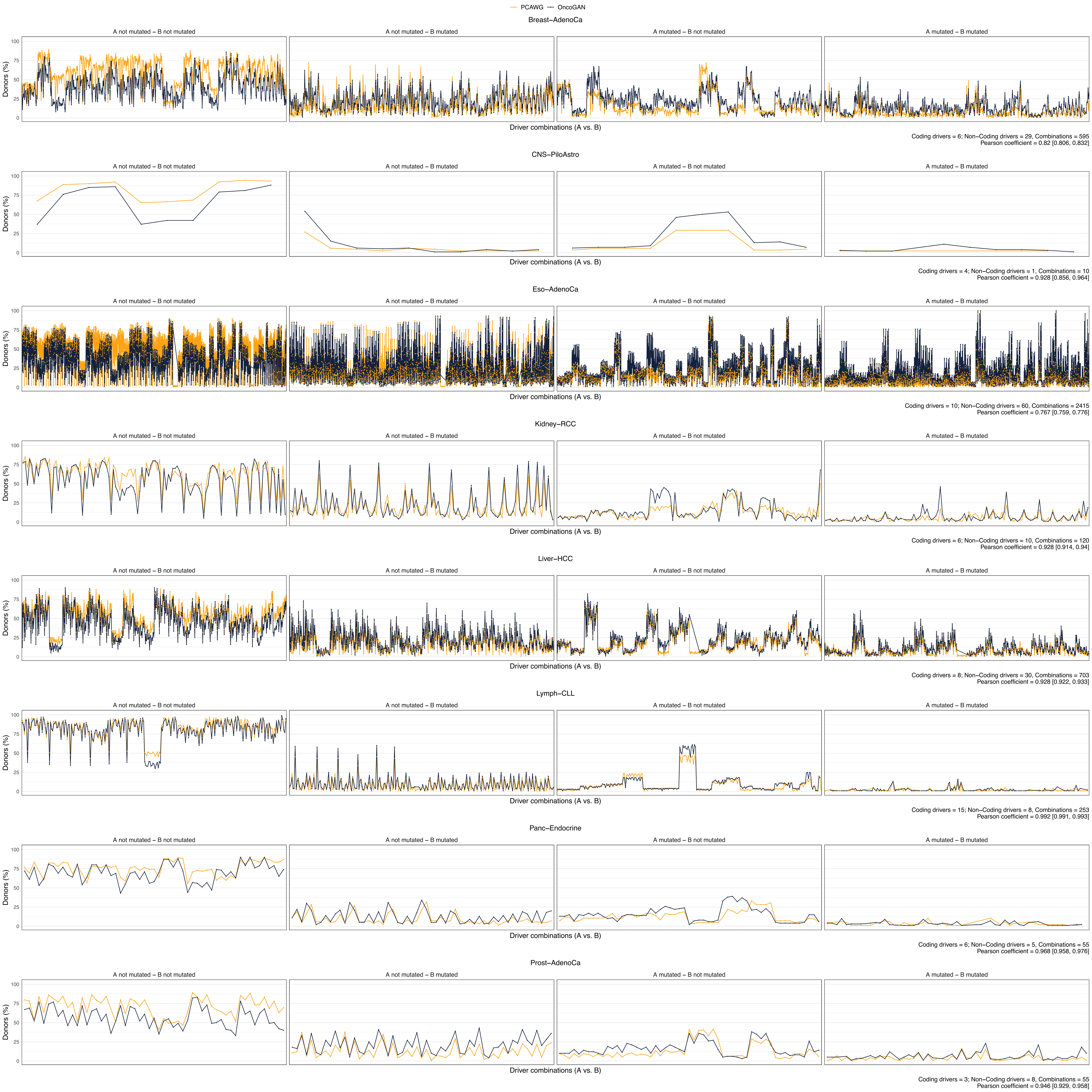
PCAWG
OncoGAN
Breast−AdenoCa
A not mutated − B not mutated
A not mutated − B mutated
A mutated − B not mutated
A mutated − B mutated
Donors (%)
Driver combinations (A vs. B)
Coding drivers = 6; Non−Coding drivers = 29, Combinations = 595
Pearson coefficient = 0.82 [0.806, 0.832]
CNS−PiloAstro
Coding drivers = 4; Non−Coding drivers = 1, Combinations = 10
Pearson coefficient = 0.928 [0.856, 0.964]
Eso−AdenoCa
Coding drivers = 10; Non−Coding drivers = 60, Combinations = 2415
Pearson coefficient = 0.767 [0.759, 0.776]
Kidney−RCC
Coding drivers = 6; Non−Coding drivers = 10, Combinations = 120
Pearson coefficient = 0.928 [0.914, 0.94]
Liver−HCC
Coding drivers = 8; Non−Coding drivers = 30, Combinations = 703
Pearson coefficient = 0.928 [0.922, 0.933]
Lymph−CLL
Coding drivers = 15; Non−Coding drivers = 8, Combinations = 253
Pearson coefficient = 0.992 [0.991, 0.993]
Panc−Endocrine
Coding drivers = 6; Non−Coding drivers = 5, Combinations = 55
Pearson coefficient = 0.968 [0.958, 0.976]
Prost−AdenoCa
Coding drivers = 3; Non−Coding drivers = 8, Combinations = 55
Pearson coefficient = 0.946 [0.929, 0.958]

**Supplementary Figure S5.** Driver correlation analysis plot for the seven tumor types. The X-axis shows all possible combinations between two driver genes, with each dot representing one 1-vs-1 combination. The Y-axis represents the percentage of donors in which that combination occurs. The number of drivers used and the total number of combinations are reported in the caption for each tumor type. The Pearson coefficient, with values exceeding 0.9 in almost all tumors, indicates a strong simulation of driver relationships. The data used to create this plot, including all possible driver combinations and their values, is listed in SuppTableS3.

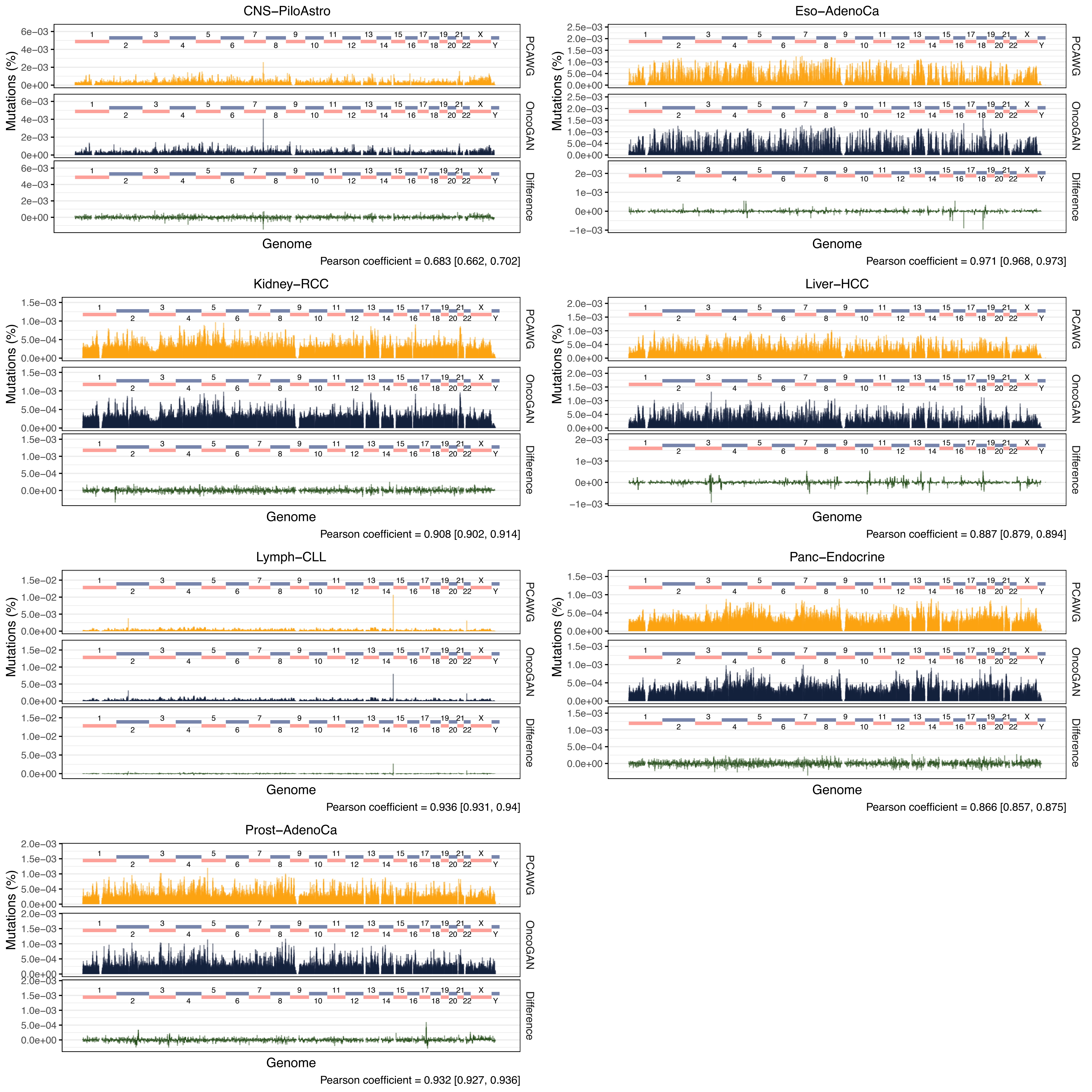

CNS−PiloAstro
Mutations (%)
PCAWG
OncoGAN
Difference
Genome
Pearson coefficient = 0.683 [0.662, 0.702]
Eso−AdenoCa
Mutations (%)
PCAWG
OncoGAN
Difference
Genome
Pearson coefficient = 0.971 [0.968, 0.973]
Kidney−RCC
Mutations (%)
PCAWG
OncoGAN
Difference
Genome
Pearson coefficient = 0.908 [0.902, 0.914]
Liver−HCC
Mutations (%)
PCAWG
OncoGAN
Difference
Genome
Pearson coefficient = 0.887 [0.879, 0.894]
Lymph−CLL
Mutations (%)
PCAWG
OncoGAN
Difference
Genome
Pearson coefficient = 0.936 [0.931, 0.94]
Panc−Endocrine
Mutations (%)
PCAWG
OncoGAN
Difference
Genome
Pearson coefficient = 0.866 [0.857, 0.875]
Prost−AdenoCa
Mutations (%)
PCAWG
OncoGAN
Difference
Genome
Pearson coefficient = 0.932 [0.927, 0.936]

**Supplementary Figure S6.** Histograms displaying the total percentage of mutations across the genome in 1Mbp bins for the remaining seven tumor types. Real donors are shown in orange, and simulated ones in black, with differences highlighted in green. Pearson correlations between PCAWG and OncoGAN genomic profiles are provided for each specific plot.

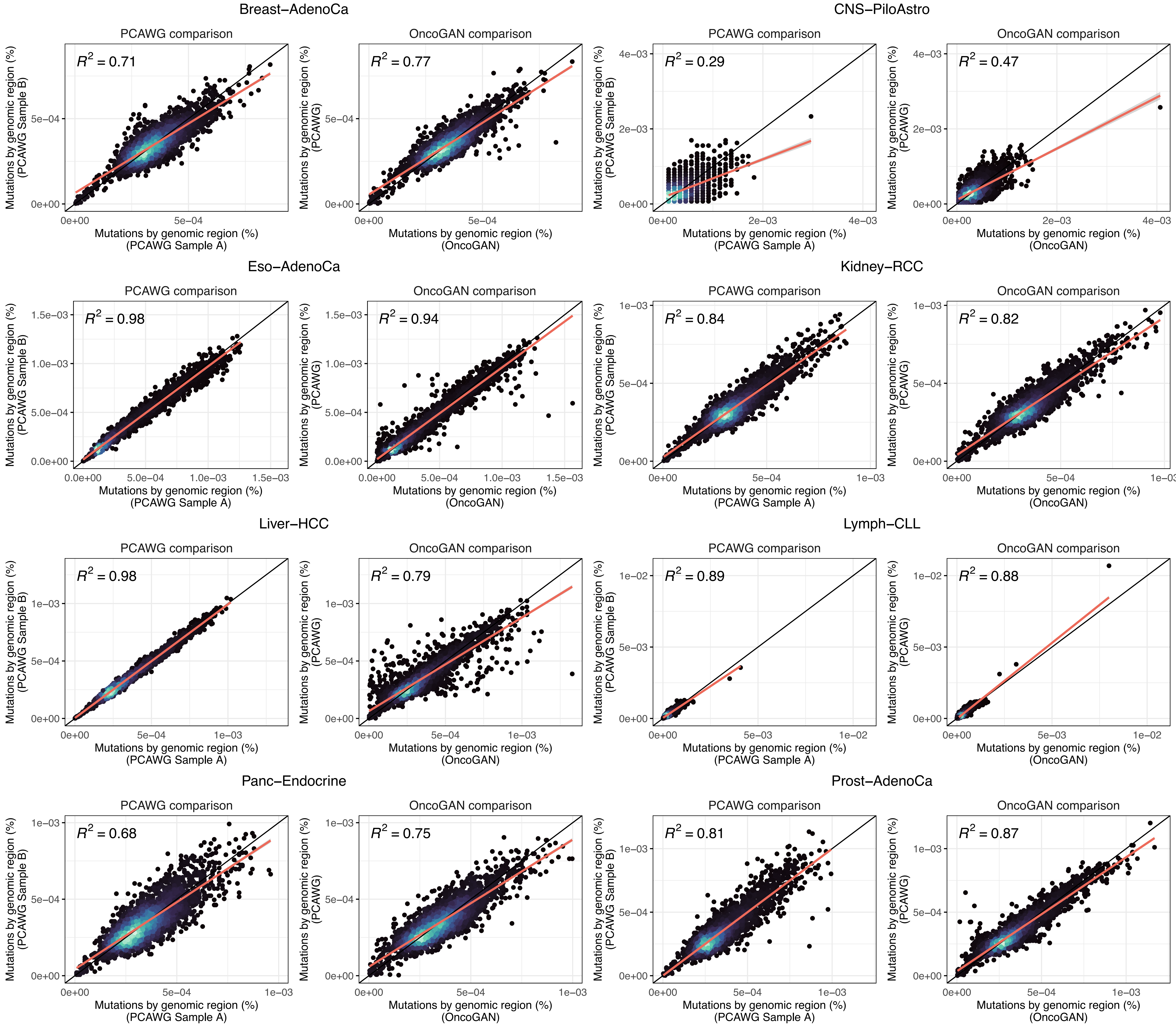

Breast−AdenoCa
CNS−PiloAstro
PCAWG comparison
OncoGAN comparison
$R^2 = 0.71$
$R^2 = 0.77$
$R^2 = 0.29$
$R^2 = 0.47$
Eso−AdenoCa
Kidney−RCC
$R^2 = 0.98$
$R^2 = 0.94$
$R^2 = 0.84$
$R^2 = 0.82$
Liver−HCC
Lymph−CLL
$R^2 = 0.98$
$R^2 = 0.79$
$R^2 = 0.89$
$R^2 = 0.88$
Panc−Endocrine
Prost−AdenoCa
$R^2 = 0.68$
$R^2 = 0.75$
$R^2 = 0.81$
$R^2 = 0.87$
Mutations by genomic region (%) (PCAWG Sample A)
Mutations by genomic region (%) (PCAWG Sample B)
Mutations by genomic region (%) (OncoGAN)
Mutations by genomic region (%) (PCAWG)

**Supplementary Figure S7.** Scatter plot comparing the percentage of mutations in each 1Mbp genomic region between two samples from the PCAWG dataset (left) and OncoGAN simulations against the entire PCAWG dataset (right) for each of the remaining tumor types. Each dot represents a region, with color indicating density; lighter colors mean higher densities. R2 values are displayed for each of the comparisons.

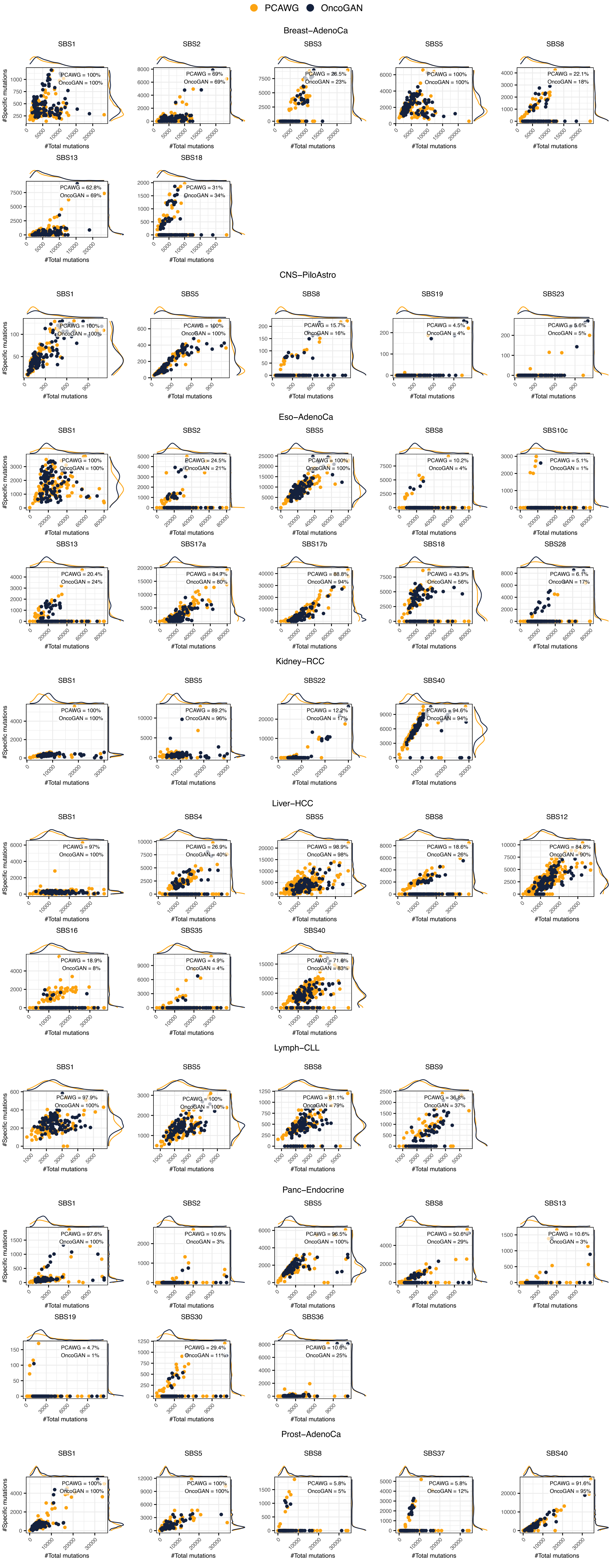

PCAWG
OncoGAN
Breast−AdenoCa
SBS1 PCAWG = 100% OncoGAN = 100%
SBS2 PCAWG = 69% OncoGAN = 69%
SBS3 PCAWG = 26.5% OncoGAN = 23%
SBS5 PCAWG = 100% OncoGAN = 100%
SBS8 PCAWG = 22.1% OncoGAN = 18%
SBS13 PCAWG = 62.8% OncoGAN = 69%
SBS18 PCAWG = 31% OncoGAN = 34%
CNS−PiloAstro
SBS1 PCAWG = 100% OncoGAN = 100%
SBS5 PCAWG = 100% OncoGAN = 100%
SBS8 PCAWG = 15.7% OncoGAN = 16%
SBS19 PCAWG = 4.5% OncoGAN = 4%
SBS23 PCAWG = 5.6% OncoGAN = 5%
Eso−AdenoCa
SBS1 PCAWG = 100% OncoGAN = 100%
SBS2 PCAWG = 24.5% OncoGAN = 21%
SBS5 PCAWG = 100% OncoGAN = 100%
SBS8 PCAWG = 10.2% OncoGAN = 4%
SBS10c PCAWG = 5.1% OncoGAN = 1%
SBS13 PCAWG = 20.4% OncoGAN = 24%
SBS17a PCAWG = 84.7% OncoGAN = 80%
SBS17b PCAWG = 88.8% OncoGAN = 94%
SBS18 PCAWG = 43.9% OncoGAN = 56%
SBS28 PCAWG = 6.1% OncoGAN = 17%
Kidney−RCC
SBS1 PCAWG = 100% OncoGAN = 100%
SBS5 PCAWG = 89.2% OncoGAN = 96%
SBS22 PCAWG = 12.2% OncoGAN = 17%
SBS40 PCAWG = 94.6% OncoGAN = 94%
Liver−HCC
SBS1 PCAWG = 97% OncoGAN = 100%
SBS4 PCAWG = 26.9% OncoGAN = 40%
SBS5 PCAWG = 98.9% OncoGAN = 98%
SBS8 PCAWG = 18.6% OncoGAN = 26%
SBS12 PCAWG = 84.8% OncoGAN = 90%
SBS16 PCAWG = 18.9% OncoGAN = 8%
SBS35 PCAWG = 4.9% OncoGAN = 4%
SBS40 PCAWG = 71.6% OncoGAN = 83%
Lymph−CLL
SBS1 PCAWG = 97.9% OncoGAN = 100%
SBS5 PCAWG = 100% OncoGAN = 100%
SBS8 PCAWG = 81.1% OncoGAN = 79%
SBS9 PCAWG = 36.8% OncoGAN = 37%
Panc−Endocrine
SBS1 PCAWG = 97.6% OncoGAN = 100%
SBS2 PCAWG = 10.6% OncoGAN = 3%
SBS5 PCAWG = 96.5% OncoGAN = 100%
SBS8 PCAWG = 50.6% OncoGAN = 29%
SBS13 PCAWG = 10.6% OncoGAN = 3%
SBS19 PCAWG = 4.7% OncoGAN = 1%
SBS30 PCAWG = 29.4% OncoGAN = 11%
SBS36 PCAWG = 10.6% OncoGAN = 25%
Prost−AdenoCa
SBS1 PCAWG = 100% OncoGAN = 100%
SBS5 PCAWG = 100% OncoGAN = 100%
SBS8 PCAWG = 5.8% OncoGAN = 5%
SBS37 PCAWG = 5.8% OncoGAN = 12%
SBS40 PCAWG = 91.6% OncoGAN = 95%
#Specific mutations
#Total mutations

**Supplementary Figure S8.** Scatter and density plots comparing the number of specific signature mutations to the total number of mutations for each donor, with real donors from PCAWG shown in orange, and simulated donors from OncoGAN shown in black. OncoGAN values correspond to those directly simulated by the tool.

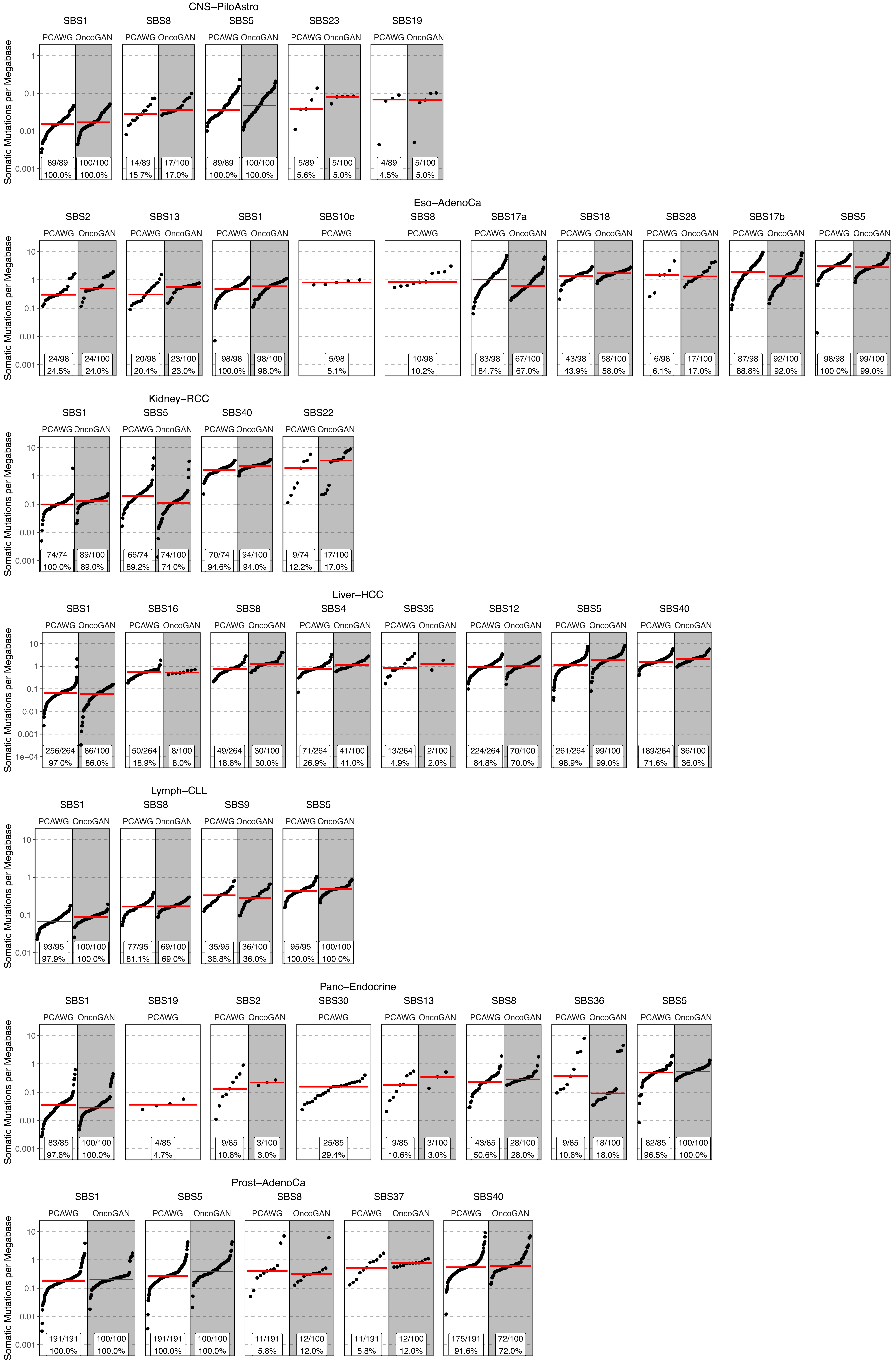
CNS−PiloAstro
SBS1 SBS8 SBS5 SBS23 SBS19
PCAWG OncoGAN
Somatic Mutations per Megabase
89/89 100.0% | 100/100 100.0% | 14/89 15.7% | 17/100 17.0% | 89/89 100.0% | 100/100 100.0% | 5/89 5.6% | 5/100 5.0% | 4/89 4.5% | 5/100 5.0%
Eso−AdenoCa
SBS2 SBS13 SBS1 SBS10c SBS8 SBS17a SBS18 SBS28 SBS17b SBS5
24/98 24.5% | 24/100 24.0% | 20/98 20.4% | 23/100 23.0% | 98/98 100.0% | 98/100 98.0% | 5/98 5.1% | 10/98 10.2% | 83/98 84.7% | 67/100 67.0% | 43/98 43.9% | 58/100 58.0% | 6/98 6.1% | 17/100 17.0% | 87/98 88.8% | 92/100 92.0% | 98/98 100.0% | 99/100 99.0%
Kidney−RCC
SBS1 SBS5 SBS40 SBS22
74/74 100.0% | 89/100 89.0% | 66/74 89.2% | 74/100 74.0% | 70/74 94.6% | 94/100 94.0% | 9/74 12.2% | 17/100 17.0%
Liver−HCC
SBS1 SBS16 SBS8 SBS4 SBS35 SBS12 SBS5 SBS40
256/264 97.0% | 86/100 86.0% | 50/264 18.9% | 8/100 8.0% | 49/264 18.6% | 30/100 30.0% | 71/264 26.9% | 41/100 41.0% | 13/264 4.9% | 2/100 2.0% | 224/264 84.8% | 70/100 70.0% | 261/264 98.9% | 99/100 99.0% | 189/264 71.6% | 36/100 36.0%
Lymph−CLL
SBS1 SBS8 SBS9 SBS5
93/95 97.9% | 100/100 100.0% | 77/95 81.1% | 69/100 69.0% | 35/95 36.8% | 36/100 36.0% | 95/95 100.0% | 100/100 100.0%
Panc−Endocrine
SBS1 SBS19 SBS2 SBS30 SBS13 SBS8 SBS36 SBS5
83/85 97.6% | 100/100 100.0% | 4/85 4.7% | 9/85 10.6% | 3/100 3.0% | 25/85 29.4% | 9/85 10.6% | 3/100 3.0% | 43/85 50.6% | 28/100 28.0% | 9/85 10.6% | 18/100 18.0% | 82/85 96.5% | 100/100 100.0%
Prost−AdenoCa
SBS1 SBS5 SBS8 SBS37 SBS40
191/191 100.0% | 100/100 100.0% | 191/191 100.0% | 100/100 100.0% | 11/191 5.8% | 12/100 12.0% | 11/191 5.8% | 12/100 12.0% | 175/191 91.6% | 72/100 72.0%
10 1 0.1 0.01 0.001 1e−04

**Supplementary Figure S9.** Mutational signatures distributions detected by SigProfiler showing the number of somatic mutations per megabase and the percentage of donors exhibiting each signature for the remaining tumor types. Dots represent individual donors; the red line illustrates the mean number of somatic mutations per megabase. *SBS, single base substitutions*.

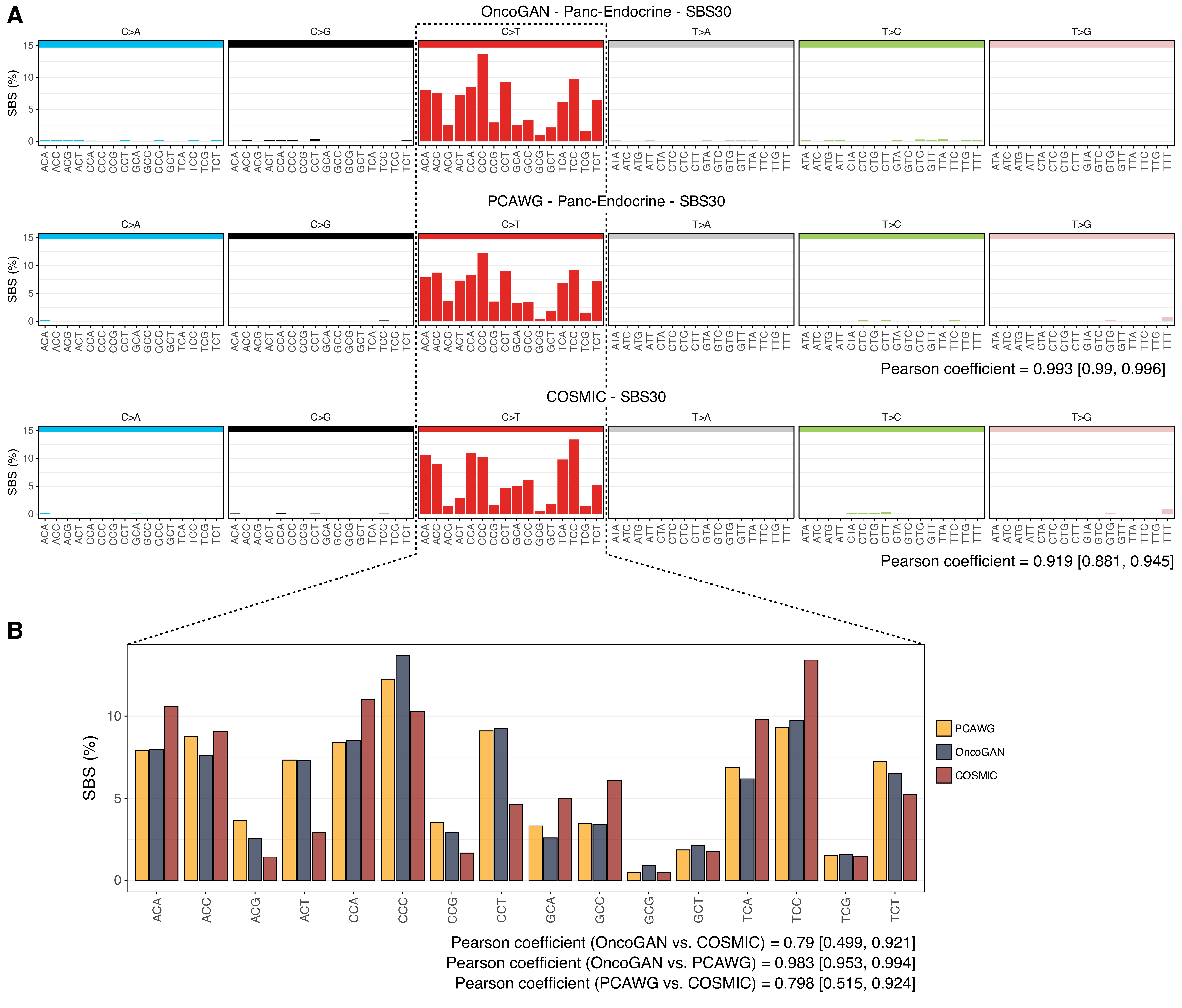
A
OncoGAN - Panc-Endocrine - SBS30
C>A
C>G
C>T
T>A
T>C
T>G
SBS (%)
PCAWG - Panc-Endocrine - SBS30
Pearson coefficient = 0.993 [0.99, 0.996]
COSMIC - SBS30
Pearson coefficient = 0.919 [0.881, 0.945]
B
SBS (%)
PCAWG
OncoGAN
COSMIC
ACA
ACC
ACG
ACT
CCA
CCC
CCG
CCT
GCA
GCC
GCG
GCT
TCA
TCC
TCG
TCT
Pearson coefficient (OncoGAN vs. COSMIC) = 0.79 [0.499, 0.921]
Pearson coefficient (OncoGAN vs. PCAWG) = 0.983 [0.953, 0.994]
Pearson coefficient (PCAWG vs. COSMIC) = 0.798 [0.515, 0.924]

**Supplementary Figure S10.** Comparison of the mutational pattern for the SBS30 signature in the Eso-AdenoCa tumor type across the PCAWG, OncoGAN, and COSMIC datasets. A) For all trinucleotide contexts the percentage each specific context contributes to the signature. B) Focusing on the C>T context, as it is the predominant altered context in this signature.

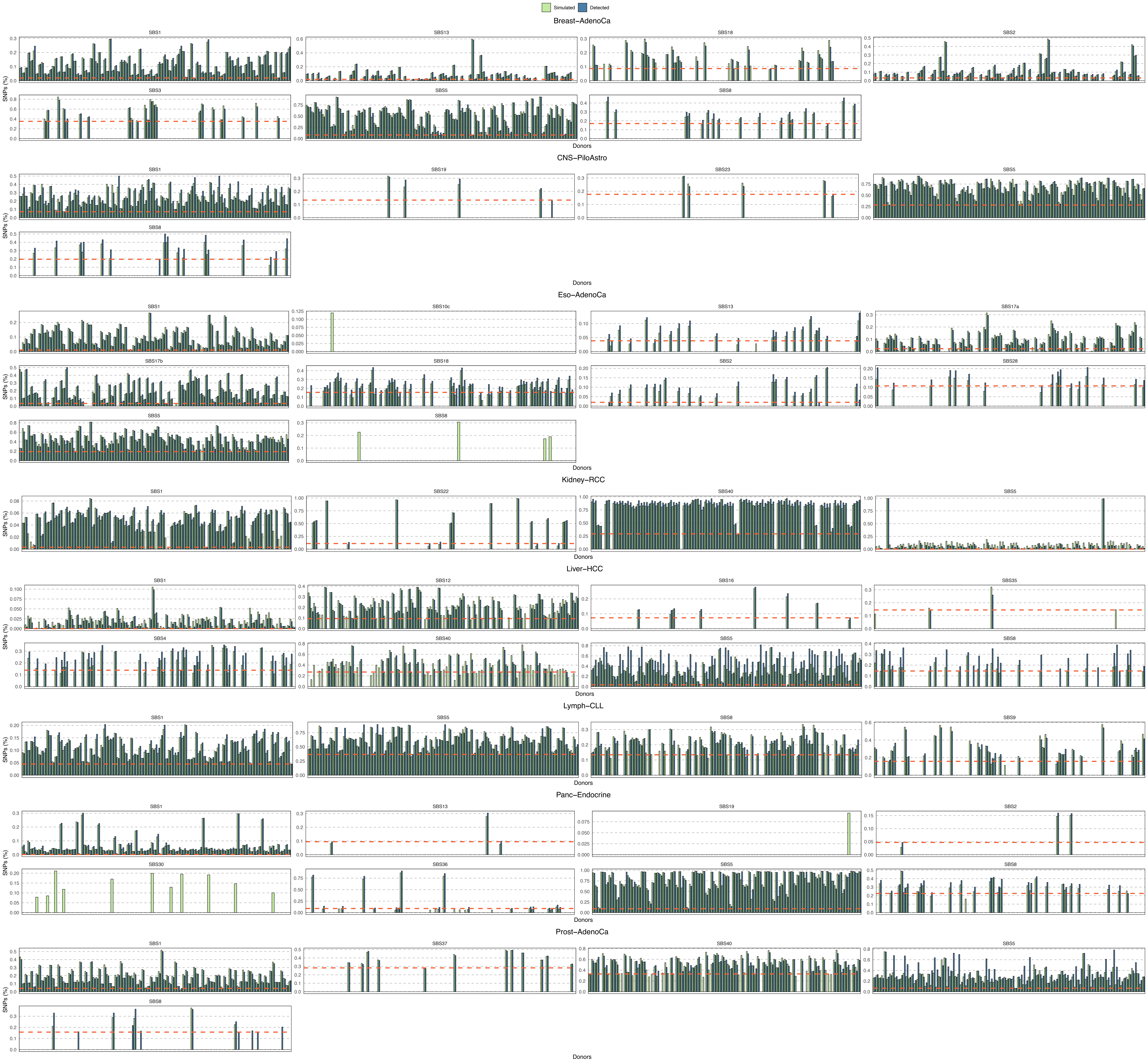

Simulated
Detected
Breast−AdenoCa
SBS1
SBS13
SBS18
SBS2
SBS3
SBS5
SBS8
SNPs (%)
Donors
CNS−PiloAstro
SBS1
SBS19
SBS23
SBS5
SBS8
SNPs (%)
Donors
Eso−AdenoCa
SBS1
SBS10c
SBS13
SBS17a
SBS17b
SBS18
SBS2
SBS28
SBS5
SBS8
SNPs (%)
Donors
Kidney−RCC
SBS1
SBS22
SBS40
SBS5
SNPs (%)
Donors
Liver−HCC
SBS1
SBS12
SBS16
SBS35
SBS4
SBS40
SBS5
SBS8
SNPs (%)
Donors
Lymph−CLL
SBS1
SBS5
SBS8
SBS9
SNPs (%)
Donors
Panc−Endocrine
SBS1
SBS13
SBS19
SBS2
SBS30
SBS36
SBS5
SBS8
SNPs (%)
Donors
Prost−AdenoCa
SBS1
SBS37
SBS40
SBS5
SBS8
SNPs (%)
Donors

**Supplementary Figure S11.** Comparison of the percentage of mutations directly simulated (OncoGAN, green) and detected (SigProfiler, blue) for each mutational signature across donors within the eight studied tumor types. Red dashed line indicates the lowest percentage of mutations detected by SigProfiler. *SNP, Single nucleotide polymorphism.*

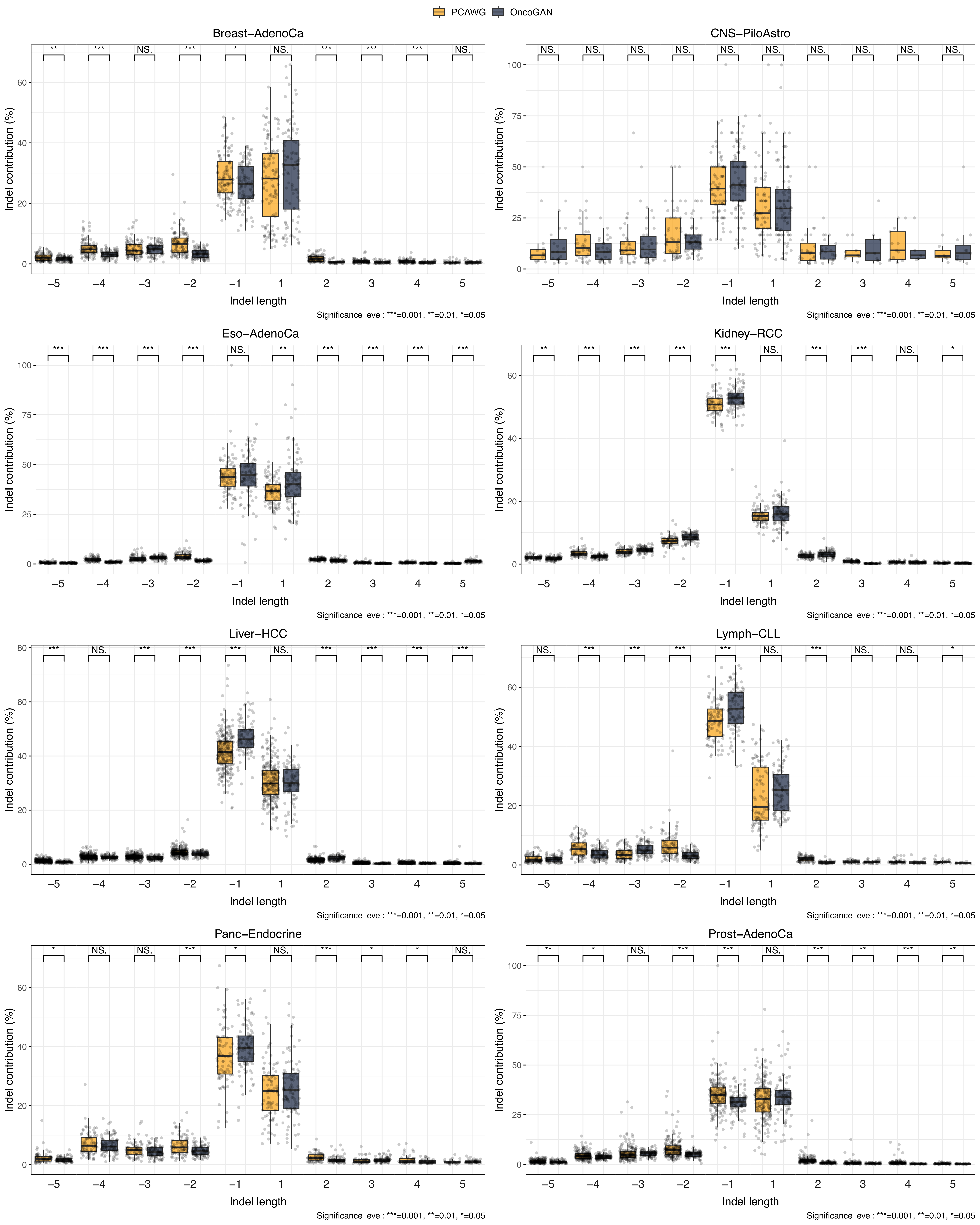
PCAWG
OncoGAN
Breast−AdenoCa
** *** NS. *** * NS. *** *** *** NS.
CNS−PiloAstro
NS. NS. NS. NS. NS. NS. NS. NS. NS. NS.
Eso−AdenoCa
*** *** *** *** NS. ** *** *** *** ***
Kidney−RCC
** *** *** *** *** NS. *** *** NS. *
Liver−HCC
*** NS. *** *** *** NS. *** *** *** ***
Lymph−CLL
NS. *** *** *** *** NS. *** NS. NS. *
Panc−Endocrine
* NS. NS. *** * NS. *** * * NS.
Prost−AdenoCa
** * NS. *** *** NS. *** ** *** **
Indel contribution (%)
Indel length
−5 −4 −3 −2 −1 1 2 3 4 5
Significance level: ***=0.001, **=0.01, *=0.05

**Supplementary Figure S12.** Boxplots comparing indel length distribution between PCAWG (orange) and OncoGAN (black). Negative X values represent deletions, and positive X values represent insertions. The Y-axis shows the contribution (%) of each specific indel length relative to the total number of indels per donor. Only indels up to a size of 5 are plotted. The Wilcoxon test was used to compare the groups. *NS.: p-value > 0.05; *: p-value <= 0.05; **: p-value <= 0.01; ***: p-value <= 0.001*

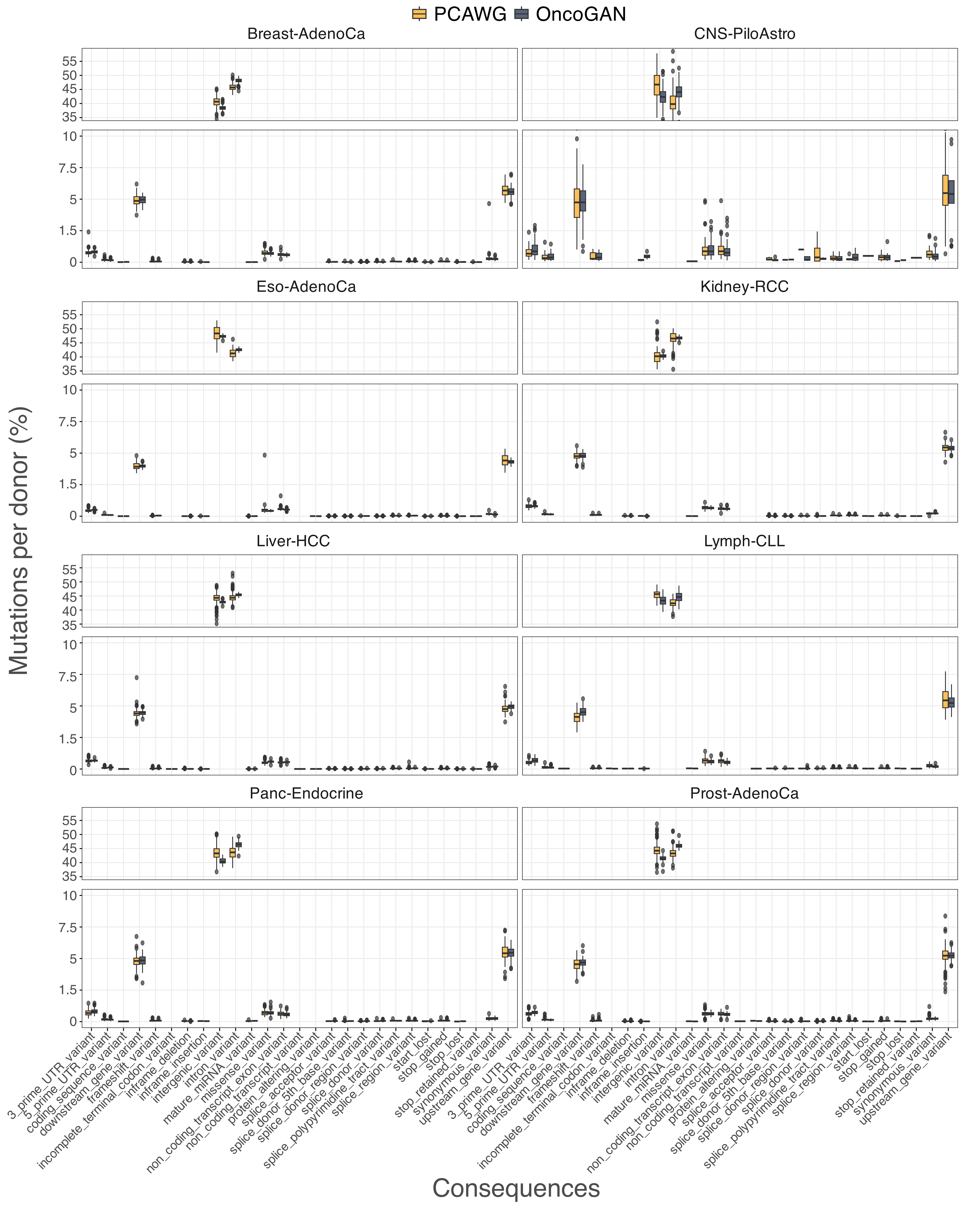

PCAWG
OncoGAN
Breast-AdenoCa
CNS-PiloAstro
Eso-AdenoCa
Kidney-RCC
Liver-HCC
Lymph-CLL
Panc-Endocrine
Prost-AdenoCa
Mutations per donor (%)
Consequences
3_prime_UTR_variant
5_prime_UTR_variant
coding_sequence_variant
downstream_gene_variant
frameshift_variant
incomplete_terminal_codon_variant
inframe_deletion
inframe_insertion
intergenic_variant
intron_variant
mature_miRNA_variant
missense_variant
non_coding_transcript_exon_variant
non_coding_transcript_variant
protein_altering_variant
splice_acceptor_variant
splice_donor_5th_base_variant
splice_donor_region_variant
splice_donor_variant
splice_polypyrimidine_tract_variant
splice_region_variant
start_lost
stop_gained
stop_lost
stop_retained_variant
synonymous_variant
upstream_gene_variant

**Supplementary Figure S13.** Boxplots comparing the frequency of predicted effects for real (orange) and simulated (black) mutations per donor using the Variant Effect Predictor (VEP) tool. The X-axis lists the possible mutation consequences considered by VEP, while the Y-axis shows the percentage of mutations corresponding to each consequence per donor.

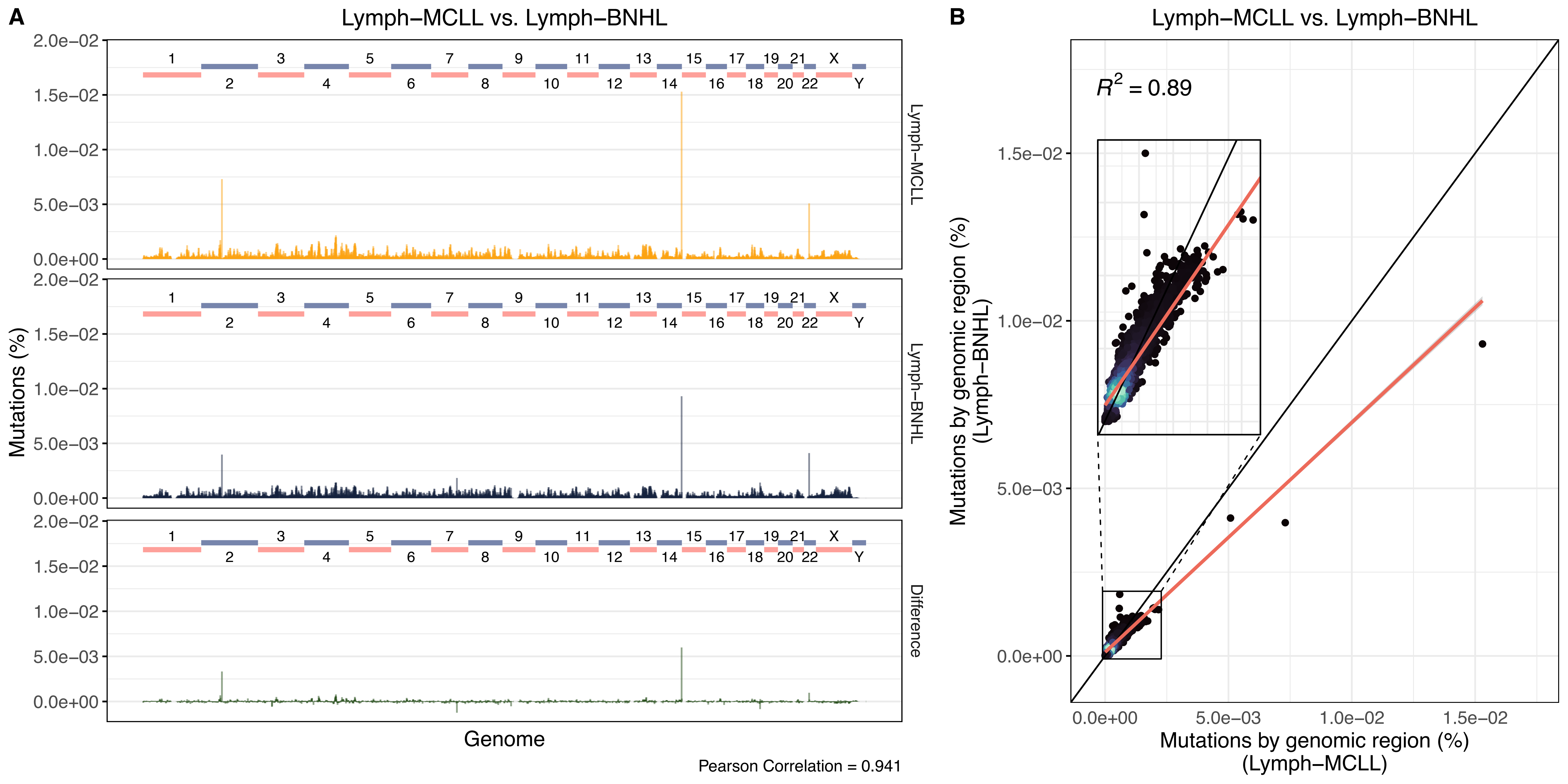

A
Lymph−MCLL vs. Lymph−BNHL
Lymph−MCLL
Lymph−BNHL
Difference
Mutations (%)
Genome
Pearson Correlation = 0.941
B
Lymph−MCLL vs. Lymph−BNHL
$R^2 = 0.89$
Mutations by genomic region (%) (Lymph−BNHL)
Mutations by genomic region (%) (Lymph−MCLL)

**Supplementary Figure S14.** Comparison of real Lymph-MCLL and Lymph-BNHL tumor types illustrating their similarity in terms of genomic mutation profiles. A) Histogram displaying the total percentage of mutations across the genome in 1Mbp bins. Real donors are shown in orange, and simulated ones in black, with differences highlighted in green. The Pearson correlation is 0.941, indicating a high degree of similarity. B) Scatter plot comparing the percentage of mutations in each 1Mbp genomic region between both tumor types. Each dot represents a region, with color indicating density; lighter colors indicate higher densities.

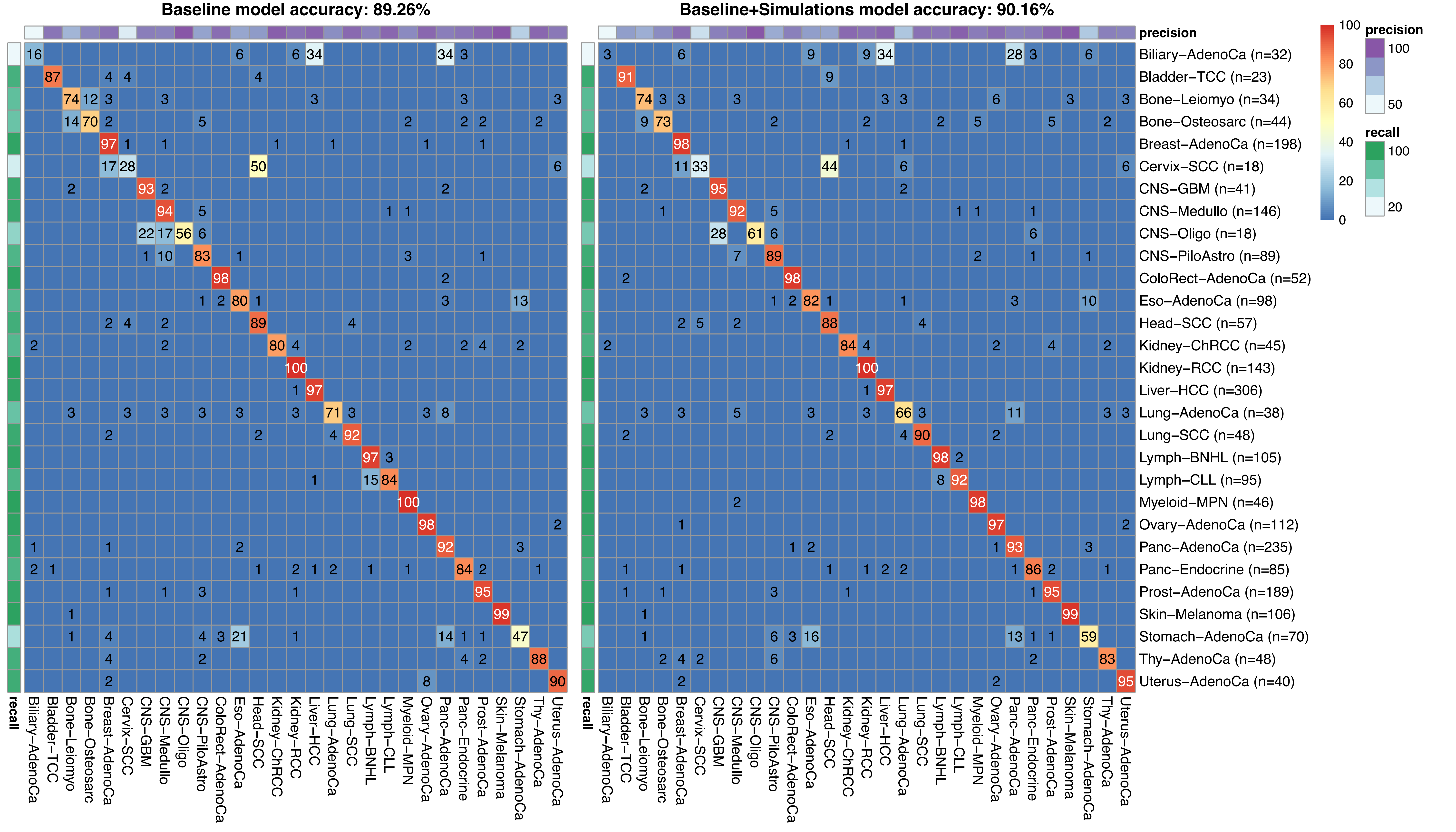

Baseline model accuracy: 89.26%
Baseline+Simulations model accuracy: 90.16%
precision
recall
100
80
60
40
20
0
precision
100
50
recall
100
20
Biliary–AdenoCa (n=32)
Bladder–TCC (n=23)
Bone–Leiomyo (n=34)
Bone–Osteosarc (n=44)
Breast–AdenoCa (n=198)
Cervix–SCC (n=18)
CNS–GBM (n=41)
CNS–Medullo (n=146)
CNS–Oligo (n=18)
CNS–PiloAstro (n=89)
ColoRect–AdenoCa (n=52)
Eso–AdenoCa (n=98)
Head–SCC (n=57)
Kidney–ChRCC (n=45)
Kidney–RCC (n=143)
Liver–HCC (n=306)
Lung–AdenoCa (n=38)
Lung–SCC (n=48)
Lymph–BNHL (n=105)
Lymph–CLL (n=95)
Myeloid–MPN (n=46)
Ovary–AdenoCa (n=112)
Panc–AdenoCa (n=235)
Panc–Endocrine (n=85)
Prost–AdenoCa (n=189)
Skin–Melanoma (n=106)
Stomach–AdenoCa (n=70)
Thy–AdenoCa (n=48)
Uterus–AdenoCa (n=40)
Biliary–AdenoCa
Bladder–TCC
Bone–Leiomyo
Bone–Osteosarc
Breast–AdenoCa
Cervix–SCC
CNS–GBM
CNS–Medullo
CNS–Oligo
CNS–PiloAstro
ColoRect–AdenoCa
Eso–AdenoCa
Head–SCC
Kidney–ChRCC
Kidney–RCC
Liver–HCC
Lung–AdenoCa
Lung–SCC
Lymph–BNHL
Lymph–CLL
Myeloid–MPN
Ovary–AdenoCa
Panc–AdenoCa
Panc–Endocrine
Prost–AdenoCa
Skin–Melanoma
Stomach–AdenoCa
Thy–AdenoCa
Uterus–AdenoCa

**Supplementary Figure S15.** DeepTumour heatmap displaying the accuracy of the baseline and the new classifier using a held-out portion of the PCAWG data set for evaluation (5-fold cross-validation). Baseline model was trained using only PCAWG samples, whereas the new model employs a mix of real and synthetic samples. Each row corresponds to the true tumor type and columns correspond to the class predictions emitted by DeepTumour. Cells are labeled with the percentage of donors of a particular type that were classified by DeepTumour as a particular type. The recall and precision of each classifier are shown in the color bars at the top and left sides of the matrix. All values represent the mean of 5 runs using selected data set partitions.

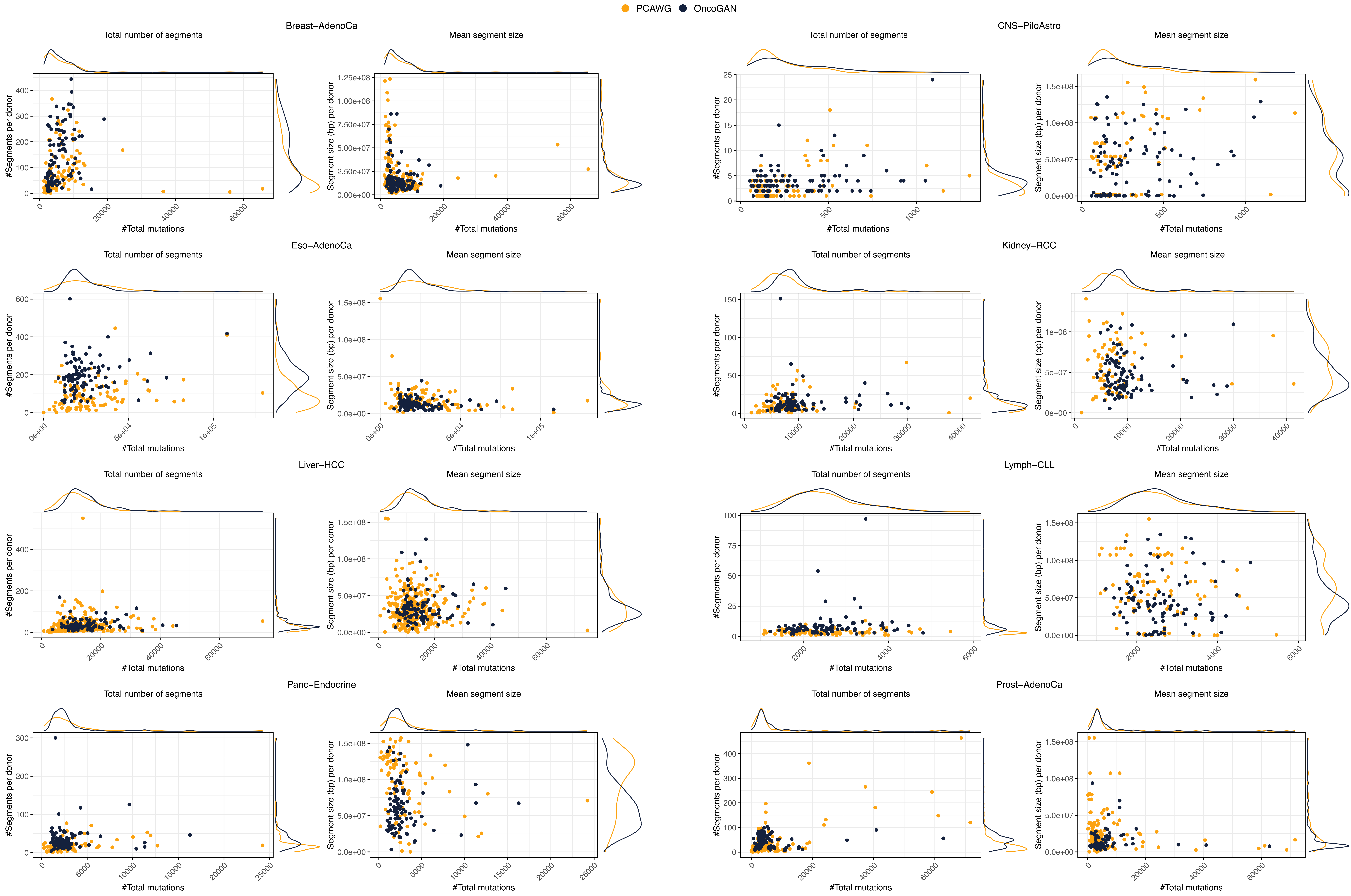

PCAWG
OncoGAN
Breast−AdenoCa
CNS−PiloAstro
Eso−AdenoCa
Kidney−RCC
Liver−HCC
Lymph−CLL
Panc−Endocrine
Prost−AdenoCa
Total number of segments
Mean segment size
#Segments per donor
Segment size (bp) per donor
#Total mutations

**Supplementary Figure S16.** Scatter and density plots showing the relationship between the total number of mutations per donor and two chromosomal instability scores: the total number of aberrant segments and their mean length. As shown, the distributions of real (orange) and simulated (black) donors are highly similar.

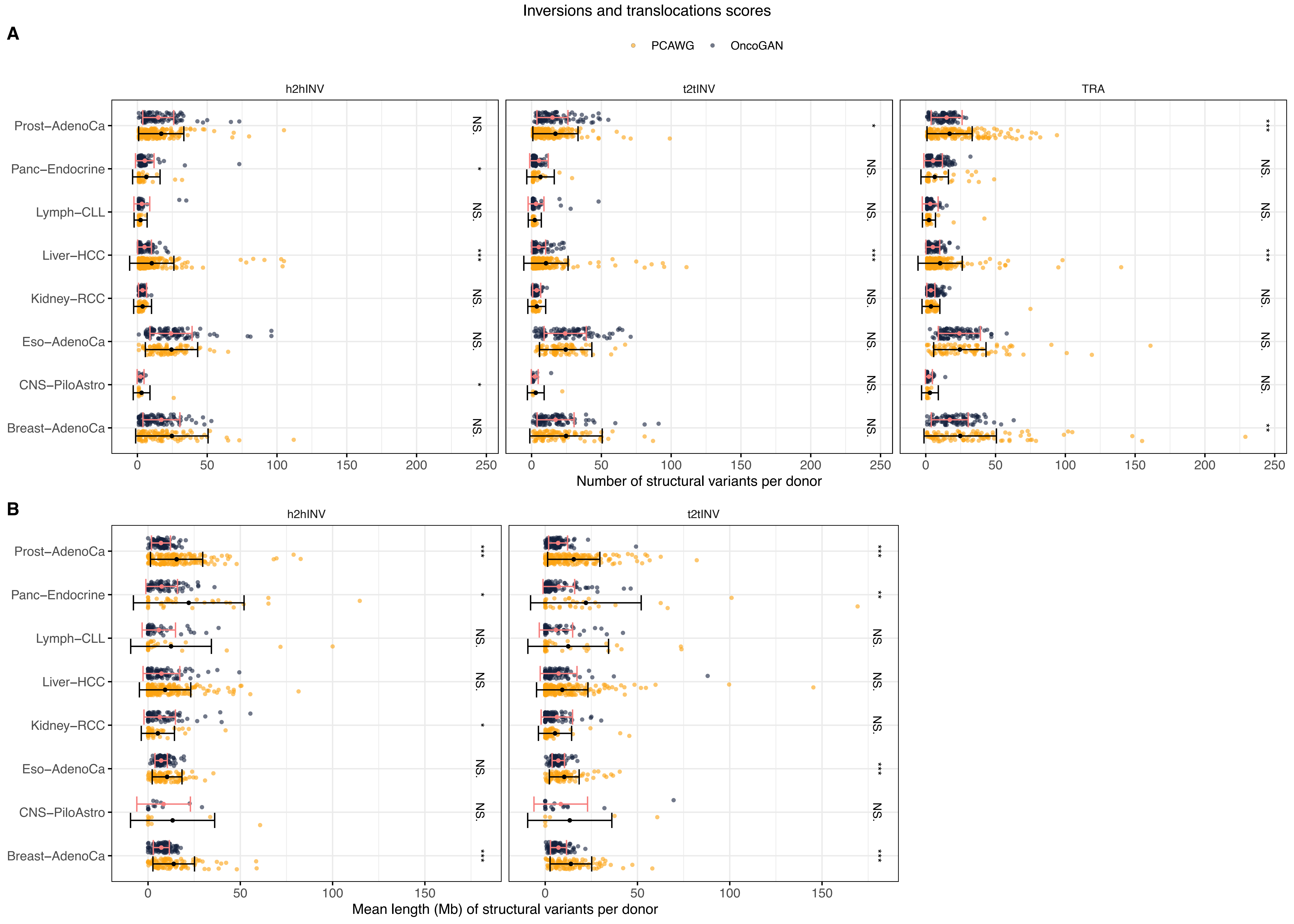
Inversions and translocations scores
A
PCAWG
OncoGAN
h2hINV
t2tINV
TRA
Prost−AdenoCa
Panc−Endocrine
Lymph−CLL
Liver−HCC
Kidney−RCC
Eso−AdenoCa
CNS−PiloAstro
Breast−AdenoCa
NS.
*
***
**
0
50
100
150
200
250
Number of structural variants per donor
B
h2hINV
t2tINV
Mean length (Mb) of structural variants per donor

**Supplementary Figure S17.** Chromosomal instability scores measuring the similarity of inversion and translocation SVs between real (orange) and simulated (black) donors. A) Number of h2hINV, t2tINV, and TRA events per donor. B) Mean length of SVs in base pairs. Error bars represent the standard deviation of the dataset. The Wilcoxon test was used to compare the groups. *NS.: p-value > 0.05; *: p-value <= 0.05; **: p-value <= 0.01; ***: p-value <= 0.001. h2hINV, head-to-head inversion; t2tINV, tail-to-tail inversion; TRA, Translocation*.
